# Behavioral, Cardiovascular, and Neuroinflammatory Effects of 3-Aminobenzamide in a Rodent Model of Chronic Psychological Stress

**DOI:** 10.64898/2026.04.28.721400

**Authors:** Liza J. Wills, Hui Wang-Heaton, Aaron J. Polichnowski, Kristy L. Thomas, Benjamin E. Jewett, Seth Jewett, Grayson Aldridge, Gregory A. Ordway, Russell W. Brown, Michelle J. Chandley

## Abstract

**Background:** Major depressive disorder (MDD) affects approximately 20% of the population, with over 30% of cases demonstrating treatment resistance. Postmortem analyses have revealed increased poly (ADP-ribose) polymerase 1 (PARP-1) expression in prefrontal cortical white matter of individuals with MDD, suggesting PARP-1 as a potential therapeutic target. Chronic stress, a major risk factor for depression, affects multiple physiological domains including behavior, cardiovascular function, neuroinflammation, and gut-brain axis signaling.

**Methods:** We conducted a comprehensive multi-system investigation of PARP inhibition effects on stress-induced pathophysiology using the social defeat stress/chronic unpredictable stress (SDS+CUS) rodent model. In the primary study, male Sprague-Dawley rats (N=32) underwent 10 days of SDS+CUS while receiving daily treatment with the PARP inhibitor 3-aminobenzamide (3-AB; 40mg/kg), selective serotonin reuptake inhibitor fluoxetine (FLX; 10mg/kg), or saline (0.9% NaCl), with non-stressed controls included. Behavioral outcomes were assessed via sucrose preference and social interaction tests. Neurobiological analyses examined PARP-1 expression, microglial morphology, and proinflammatory cytokine levels (IL-1β, TNF-α, IL-6) in relevant brain regions. In a parallel cardiovascular study, a separate cohort of stressed rats (N=8) received either saline or 3-AB treatment while hemodynamic parameters were monitored via telemetry before, during, and after stress exposure.

**Results:** Saline-treated stressed rats demonstrated significantly reduced sucrose preference compared to the 3-AB/Stress rats and social avoidance compared to the 3-AB/Stress and Control/ No-Stress groups. Cardiovascular monitoring showed increases in systolic and mean arterial pressure and a decrease in heart rate across the paradigm; Treatment x Time interations for blood pressure were not significant, while heart rate decline was greater in saline-treated rats. Neurobiological analyses of PARP-1 expression in PFC did not reveal significant group differences. Microglial morphological analysis revealed a shift toward more prolate soma morphology in the saline-treated stressed rats compared to the non-stressed controls. Additionally, 3-AB-treated rats showed significantly more oblate microglia than saline-treated rats. Saline-treated stressed rats exhibited significantly increased hippocampal proinflammatory cytokines, with 3-AB treatment specifically attenuating TNF-α and IL-1β levels.

**Conclusion:** 3-AB treatment provided multi-system protection against chronic stress effects, preventing behavioral deficits and attenuating select markers of neuroinflammation, without significantly altering blood pressure. These findings suggest that 3-AB, and potentially PARP-1 inhibition, warrants further investigation as a therapeutic approach for stress-related disorders.

**Significant Outcomes:**

- 3-aminobenzamide (3-AB) treatment was associated with reduced anhedonia-like behavior and social avoidance relative to saline-treated stressed rats in a validated rodent model combining social defeat and chronic unpredictable stress.
- 3-AB did not significantly alter stress-related changes in blood pressure; heart rate was nominally greater in saline-treated rats.
- 3-AB treatment lead to more oblate microglial morphology in PFC gray matter and significantly attenuated the stress-induced increase in hippocampal TNF-α and IL-1β.
- The multi-domain protective effects of 3-AB, spanning behavioral and neuroinflammatory measures, support further investigation of this compound as a potential therapeutic strategy for stress-related disorders.

**Limitations:**

- This study examined only male rats, limiting generalizability to female subjects despite the higher prevalence of MDD in women.
- Behavioral assessments were limited to sucrose preference and social interaction tests; additional tests of other depression-relevant behaviors would provide a more comprehensive phenotypic profile.
- We did not find changes of PARP-1 expression in the PFC, other brain regions should be examined in the future.
- Long-term effects of 3-AB treatment beyond the 10-day stress paradigm remain unexplored.

## Introduction

Major depressive disorder (MDD) is a leading cause of disability worldwide, with a lifetime prevalence rate of approximately 20%^1,2^. Despite the prevalence and impact of MDD, current pharmacological treatments remain inadequate, with over 30% of patients demonstrating treatment resistance^3,4^. This substantial treatment gap highlights the critical need for novel antidepressant approaches based on improved understanding of MDD pathophysiology beyond monoaminergic mechanisms.

Converging evidence indicates that neuroinflammation plays a significant role in MDD etiology^5,6^. Individuals with MDD exhibit elevated levels of inflammatory markers, including proinflammatory cytokines such as interleukin-1β (IL-1β), interleukin-6 (IL-6), and tumor necrosis factor-alpha (TNF-α) in key regions known for mood regulation such as the prefrontal cortex (PFC) and hippocampus (HPC)^7^ . Additionally, the bidirectional relationship between peripheral inflammation and central neuroinflammatory processes is now well-established, with several pathways identified for immune-brain communication^8^ . MDD frequently co-occurs with conditions characterized by chronic inflammation, suggesting common underlying pathological mechanisms^9,10^ .

Chronic psychological stress, a major risk factor for depression, is associated with increased reactive oxygen species (ROS) production, oxidative stress, and subsequent deoxyribonucleic acid (DNA) damage^11^ . Microglia, the resident immune cells of the central nervous system (CNS), are particularly sensitive to stress signals and can become primed or activated during chronic stress, releasing proinflammatory cytokines and ROS that further exacerbate neuroinflammation^12–14^.

This creates a potential positive feedback loop between oxidative stress, microglial activation, and neuroinflammation that may contribute to the development and maintenance of depressive symptomatology.

The systemic nature of chronic stress extends beyond neurobiological changes to encompass cardiovascular dysfunction. Chronic stress exposure is associated with hypertension, altered heart rate variability, and elevated cardiovascular disease risk^15–17^. These multi-system effects suggest that effective antidepressant interventions may need to address not only central neurobiological mechanisms but also peripheral physiological dysfunction.

Poly (ADP-ribose) polymerase 1 (PARP-1), a nuclear enzyme primarily involved in DNA repair, has been implicated in this pathophysiological cascade. Under conditions of excessive oxidative stress, PARP-1 becomes overactivated, depleting cellular nicotinamide adenine dinucleotide (NAD^+^) reserves and promoting proinflammatory signaling through nuclear factor kappa B (NF-κB) co-activation^18,19^. Importantly, we have previously demonstrated that postmortem brain tissue from individuals with MDD have increased PARP-1 messenger ribose nucleic acid (mRNA) expression in PFC white matter compared to psychiatrically normal tissue^20^ . This suggests that PARP-1 overactivation may be a key molecular link between chronic stress, neuroinflammation, and depression.

PARP inhibitors, already U.S. Food and Drug Administration (FDA)-approved for cancer treatment, have demonstrated promise in reducing neuroinflammation in various neurological conditions^21,22^. Previous research showed that the PARP inhibitor 3-aminobenzamide (3-AB) counteracted lipopolysaccharide-induced depressive-like behaviors in mice through anti-inflammatory mechanisms^10^. Additionally, our laboratory demonstrated antidepressant-like effects of 3-AB in the forced swim test and in a combined social defeat stress/chronic unpredictable stress (SDS+CUS) paradigm^23^. More recently, a case report from our group documented rapid antidepressant and anxiolytic effects following administration of the PARP inhibitor niraparib in a patient with treatment-resistant depression^24^ . These findings collectively suggest PARP inhibition may represent a novel antidepressant strategy targeting neuroinflammatory processes.

The present study aimed to comprehensively investigate the multi-system effects of 3-AB in the SDS+CUS rodent model of depression, examining behavioral, cardiovascular, and neuroinflammatory markers. We hypothesized that 3-AB would prevent stress-induced anhedonia and social avoidance while reducing neuroinflammatory markers, prevent stress-induced increases in arterial blood pressure, and preserve gut microbiome homeostasis. Through parallel behavioral and cardiovascular studies, combined with detailed neurobiological analyses, we sought to characterize the multi-system effects of 3-AB treatment on stress-induced behavioral, cardiovascular, and neuroinflammatory outcomes, and to generate hypotheses regarding the potential involvement of PARP-1 in these effects.

## Materials and Methods

### Laboratory Animals

Adult male Sprague Dawley rats (Envigo, Indianapolis, IN), approximately 60 days old upon arrival, were used as experimental subjects (“intruders”) or controls across two cohorts: a primary behavioral/molecular cohort (n = 32) and a separate cardiovascular monitoring cohort (n = 8). These samples sizes were selected based on group sizes used in our previously published SDS+CUS protocol^23^. No formal randomization procedure was used to allocate subjects to experimental groups. No animals were excluded in the behavioral analyses. Male Long-Evans Hooded rats were selected as aggressors (“residents”) in the social defeat paradigm given their well-documented territorial aggression relative to Sprague-Dawley rats^25–27^. Aggressive behavior was confirmed through daily observation, with all resident rats exhibiting consistent attack behavior towards intruders throughout the study. Resident rats were pair-housed with ovariectomized female Sprague-Dawley rats, which were rotated daily to help maintain resident territoriality and aggression^28^. Non-stress control rats were group-housed, while stressed rats were singly housed with environmental enrichment according to National Institutes of Health (NIH) guidelines for the Care and Use of Animals. All animals were maintained on a 12h light/dark cycle with food and water available *ad libitum*. All procedures were approved by the East Tennessee State University Animal Care and Use Committee.

### Drugs and Solutions

3-Aminobenzamide (3-AB; 40mg/kg/day; CAS #: 3544-24-9) and fluoxetine HCl (FLX; 10mg/kg/day; CAS #: 56296-78-7) were obtained from Sigma-Aldrich (St. Louis, MO) and diluted in saline solution (0.9% sodium chloride). The 40mg/kg 3-AB dose was selected based on prior dose-response work from our laboratory using forced swim test, in which 3-AB doses of 0.4 and 4 mg/kg did not significantly reduce immobility or latency to immobility relative to vehicle whereas 40mg/kg produced significant antidepressant-like effects on both measures^23^. Drugs were administered daily in the morning, with 3-AB and SAL administered subcutaneously and FLX administered intraperitoneally throughout the 10-day stress protocol. FLX was included as a clinically used pharmacological comparator rather than as a validated positive control for this specific paradigm.

### Experimental Protocol

The stress model combined social defeat stress (SDS) and chronic unpredictable stress (CUS) over 10 consecutive days, as previously described and validated^20,23^. While CUS protocols conventionally employ longer durations (e.g., 14 days) when used in isolation^29,30^, our model combined paradigm layers daily social defeat on top of rotating CUS stressors, resulting in a greater cumulative stress burden per day. This combined 10-day protocol has been validated in our prior work^20,23^ to reliably produce anhedonic and social avoidance phenotypes comparable to those seen with longer single-stressor paradigms. This double-stressor design was originally adopted to reduce the likelihood of stress resilience that can develop when rodents are exposed to a single, predictable stressor administered at a consistent time each day. The SDS paradigm was conducted every morning, in which each “intruder” rat was placed in a “resident” cage for 5 minutes or until the resident performed a 2-second pin. Behaviors observed included defensive posturing, freezing, and vocal distress from intruders, and investigative sniffing, piloerection, and attack behaviors from residents. The CUS protocol exposed rats to various mild stressors at unpredictable times, including 30-min restraint, 1h shaking and crowding, 10-min cold water (18°C) swim, 15-min warm water (25°C) swim, and 24h tipped cage. Each stressor was administered twice over the 10-day period at various times in the afternoon and evening.

Animals were divided into 4 experimental groups: (1) SAL/Stress: stressed rats receiving saline (n = 8); (2) 3-AB/Stress: stressed rats receiving 3-AB (n = 8); (3) FLX/Stress: stressed rats receiving fluoxetine (n = 8); and (4) CTRL/No-Stress: non-stress control rats (n = 8).

### Assessment of Arterial Blood Pressure and Heart Rate

A separate cohort of male Sprague Dawley rats (n = 8) was used specifically for cardiovascular monitoring during the stress paradigm. This sample size is consistent with established practice in cardiovascular telemetry studies^31,32^. Animals were divided into two groups: SAL/Stress (n = 4) and 3-AB/Stress (n = 4). Acetaminophen (200mg/kg, PO; CAS #: 103-90-2) was added to animals’ water bottles 48 h prior to surgery and remained on home cages for 72 h post-operation as an analgesic. Radio-telemetric transmitters (HD-S10; Data Sciences International, St. Paul, MN) were used to assess arterial blood pressure and heart rate (HR). Under isoflurane anesthesia (2-3%, INH; CAS #: 26675-46-7), the blood pressure catheter was inserted into the right femoral artery via an inguinal incision and advanced to the abdominal aorta. The body of the transmitter was positioned intraabdominally via a midline incision and secured to the abdominal wall with sutures. Once the incision was closed, bupivacaine (TOP) was applied to the site as a local anesthetic. Rats recovered for a minimum of 7 days prior to baseline data collection. Hemodynamic parameters including systolic blood pressure (SBP), diastolic blood pressure (DBP), mean arterial pressure (MAP), and heart rate (HR) were obtained at a sampling rate of 500 Hz using Ponemah software (Data Sciences International). Baseline recordings were collected continuously during the dark cycle (1900h-0700h) for 3 consecutive days. During the stress protocol, hemodynamic data were recorded continuously during the 5-min social defeat session and again during the animal’s dark cycle. Post-stress recordings continued during the dark cycle for 3 consecutive days following protocol completion. For statistical analysis, SBP, DBP, MAP, and HR values were averaged across the 3-day baseline period and across the 3-day post-stress period separately for each animal, yielding a single baseline value and a single post-stress value per animal (n = 4/group). These per-animal averages were used for all analyses reported in the Results. The change from baseline (post-stress – baseline) was also calculated for each animal.

### Behavioral Assessments

#### Sucrose Preference Test

The sucrose preference test was conducted on experimental days 8-10 during the first 2 hours of the dark cycle (1900h – 2100h) based on a previously established paradigm^33^. Rats were presented with two pre-weighed bottles, one containing water and the other containing 0.8% (w/v) sucrose solution. Sucrose preference was calculated as the percentage of sucrose solution consumed relative to total fluid consumption. This testing window was selected to be consistent with our previously established SDS+CUS protocol^23^, though we note that because testing occurred during the final days of the stress paradigm rather than following a stress-free interval, the resulting anhedonia measure may reflect both cumulative and acute components of the stress response.

#### Social Interaction Test

The social interaction test was conducted on experimental day 11 in a locomotor arena (91cm^3^ black plexiglass box) divided by wire mesh. The experimental rat was placed on one side of the arena for a 10-min habituation period, after which a novel aggressor rat was placed on the opposite side for a 5-min interaction test. Time spent in the interaction zone (near the wire mesh) versus the avoidance zone was recorded using ANY-maze video tracking software (Stoelting Co., Wood Dale, IL).

### Tissue Collection and Processing

Animals were euthanized on experimental day 12 via live decapitation (no anesthesia). Brains were removed and divided for multiple analyses. Following sagittal bisection along the midline, the anterior left hemisphere (PFC, anterior to the dorsal HPC at approximately Bregma +1.0mm) was fixed in 4% paraformaldehyde for 24h followed by cryoprotection in 20% sucrose solution for ionized calcium-binding adaptor molecule 1 (IBA-1) immunohistochemistry (IHC). The remaining brain tissue (posterior left hemisphere containing HPC and the entire right hemisphere) was flash-frozen on dry ice and stored at - 80 for PARP-1 immunohistochemistry and cytokine analyses. Tissue was coronally sectioned using a Leica CM1950 cryostat (Buffalo Grove, IL).

### Immunohistochemistry

#### PARP-1 Immunohistochemistry

Frozen tissue sections (20µm) from the PFC were mounted on slides and fixed in cold acetone. Sections were incubated with anti-PARP-1 polyclonal antibody (1:500; Abcam Cat# ab227244, RRID: AB_3676687) overnight at 4°C, followed by donkey anti-rabbit Alexa Fluor-594 secondary antibody (1:1000; Thermo Fisher Scientific Cat# R37119, RRID: AB_2556547) for 2 hours. Images were captured using an EVOS FL Auto Imaging System (ThermoFisher Scientific) and analyzed with MCID Analysis Software (InterFocus Ltd). Three distinct regions were examined: layer 1 (L1) gray matter, layers 3-6 (L3-6) gray matter, and forceps minor of the corpus callosum (FMI) white matter. A small number of animals were excluded from specific regions due to tissue damage or unusable sections, resulting in group sizes ranging from n = 5-8 across regions (See Results).

#### IBA-1 Immunohistochemistry

Fixed tissue was sectioned (50µm) and processed as free-floating sections. Tissue was incubated with IBA-1 polyclonal antibody (1:1000; FUJIFILM Wako Pure Chemical Corporation Cat# 019-19741, RRID: AB_839504) overnight at 4°C, followed by donkey anti-rabbit Alexa Fluor-488 secondary antibody (1:1000; Thermo Fisher Scientific Cat# R37118, RRID: AB_2556546). Images were captured using a Leica TCS SP8 confocal microscope and analyzed with Imaris software (Oxford Instruments) to assess microglial morphology. Ten parameters were analyzed: soma volume, soma area, soma sphericity, ellipticity (prolate), ellipticity (oblate), filament volume, filament area, filament length, dendrite branch points, and sholl intersections.

#### Cytokine ELISAs

Ventral hippocampal tissue was collected and analyzed for proinflammatory cytokines using ELISA kits for IL-1β, TNF-α, and IL-6 (All from Biomatik Cat# IL-1β: EKF57939, TNF-α: EKF57956, IL-6: EKF57855). Tissue was homogenized in RIPA buffer with protease and phosphatase inhibitors. The protocol provided by the vendor was strictly followed. Optical density was measured at 450nm using a Bio-Tek ELx880 microplate reader and converted to protein concentration (pg/mL) using standard curves. One SAL/Stress sample was unavailable for IL-6 analysis (n = 7 for this group); all other cytokine analyses included n = 8/group.

#### Statistical Analyses

Animals were identified by numeric ID throughout data collection and analysis. Sucrose preference bottles were weighed in numerical order by animal ID, and social interaction behavior was scored automatically using ANY-maze video tracking software. In both cases, personnel performing data collection did not reference treatment group assignment. Histological (PARP-1 and IBA-1 immunohistochemistry) and ELISA analyses were similarly conducted using coded animal IDs, with the analyst not referencing treatment group assignment during quantification. Formal unblinding did not occur until after all measurements were completed. Analyses were performed using GraphPad Prism version 10.6.1 or R Studio version 2021.09.0. Normality of residuals and homogeneity of variance were assessed for all analyses prior to selecting the appropriate approach. Brown-Forsythe testing confirmed homogeneity of variance for social interaction and cytokine (TNF-⍰, IL-1β, IL-6) tests, for which ordinary or repeated-measures ANOVA with Dunnett’s post-hoc testing (or paired/unpaired t-tests, as specified below) was used. For PARP-1 IHC, Brown-Forsythe testing indicated significant variance heterogeneity across treatment groups in all three regions (L1, L3-6, FMI); Welch’s ANOVA with Dunnett’s T3 post-hoc was used for these analyses accordingly. For hemodynamic parameters, normality within each group (Shapiro-Wilk) and homogeneity of variance between groups (F-test) were assessed for baseline, post-stress, and change-from-baseline values. SBP, DBP, MAP, and HR were each analyzed by two-way repeated measures ANOVA (Treatment x Time). The Treatment x Time interaction was the primary test of whether 3-AB altered the change in each measure relative to SAL/Stress and was confirmed with unpaired t-tests on change-from-baseline values, reported as the between-group difference in change with its 95% confidence interval. Given the small sample size (n = 4/group). These analyses have limited power to detech modest effects. No correction for multiple comparisons was applied across the four hemodynamic measures, and the results are interpreted accordingly. One-way or two-way ANOVAs were used for social interaction, and cytokine analyses followed by Dunnett’s post-hoc tests comparing treatment groups against the SAL/Stress group. For the sucrose preference test, sucrose preference values were analyzed using a linear mixed-effects model with Treatment, Day, and their interaction as fixed effects and Animal ID included as a random intercept to account for repeated measurements within each animal. Post-hoc comparisons of each treatment group against the SAL/Stress group averaged across days were conducted using estimated marginal means with Šídák correction for multiple comparisons. For PARP-1 IHC, Brown-Forsythe tests revealed significant variance heterogeneity across treatment groups in all three regions examined; therefore, Welch’s ANOVA was used for each region (L1, L3-6, FMI), followed by Dunnett’s T3 post-hoc test comparing treatment groups to SAL/Stress. For microglia morphology analyses, hierarchical linear mixed-effects models were used to account for the nested structure of the data (multiple cells per animal), with treatment group as a fixed effect and Animal ID as a random effect. This approach accounts for non-independence of multiple measurements from the same animals. Post-hoc comparisons were conducted using Holm-Šídák’s multiple comparisons test. Outliers were determined using Grubbs’ test for all datasets, no data points were excluded on this basis in any analysis. Statistical significance was set at p < 0.05. Results are presented as mean ± SEM.

## Results

### Sucrose Preference Test

A linear mixed-effects model with Animal ID as a random intercept revealed significant main effects of Treatment [F(3,28) = 3.07, p = 0.044] and Day [F(2,56) = 9.77, p < 0.001] on sucrose preference, with no significant Treatment x Day interaction. Post-hoc comparisons of estimated marginal means, averaged across days, showed that the 3-AB/Stress group exhibited significantly higher sucrose preference than Sal/ Stress group (p = 0.032). The CTRL/No-Stress group and FLX/ Stress groups did not differ significantly from the SAL/Stress group (Fig 2).

**Figure 1.**
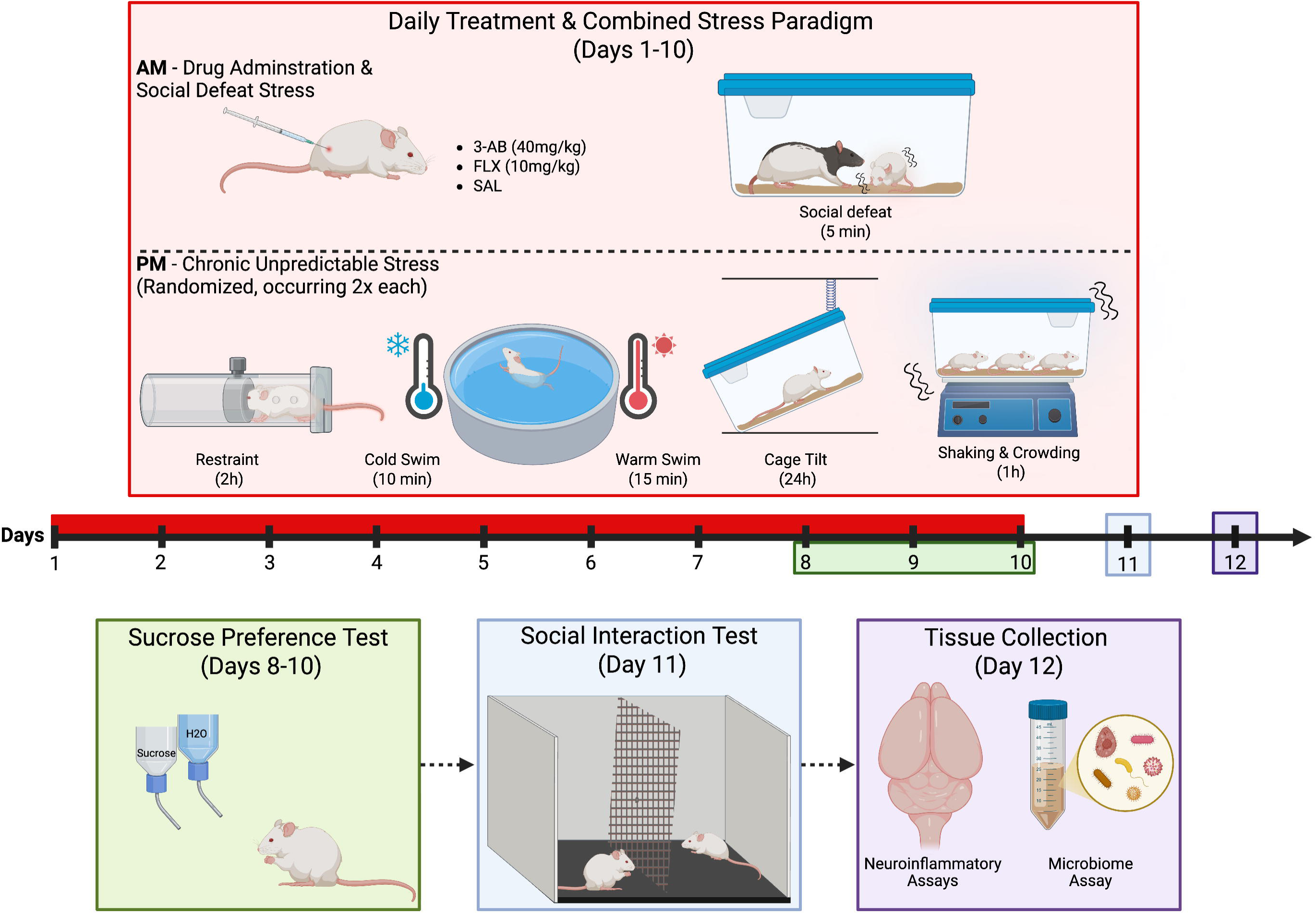
Experimental timeline and Behavioral Design. Adult Male Sprague-Dawley rats (n = 8/group) were exposed to 10 days of combined social defeat stress (SDS, 5min/day, AM) and chronic unpredictable stress (CUS, rotating PM stressors) while receiving daily injections of 3-AB (40mg/kg, s.c.), FLX (10 mg/kg, i.p.), or SAL (i.p.). CUS stressors included restraint stress (2h), cold swim (18□, 10 min), warm swim (25□, 15 min), cage tilt (24h), and shaking and crowding (1h). Anhedonia was assessed via sucrose preference testing (Days 8-10, 7PM-9PM), and social avoidance was measured on Day 11. Tissue was collected 48h post-stress (Day 12) for PARP-1 analysis (IHC), microglial morphology assessment (IHC), cytokine quantification (ELISA), and microbiome profiling (SCFA assay). Figure created using BioRender.

**Figure 2.**
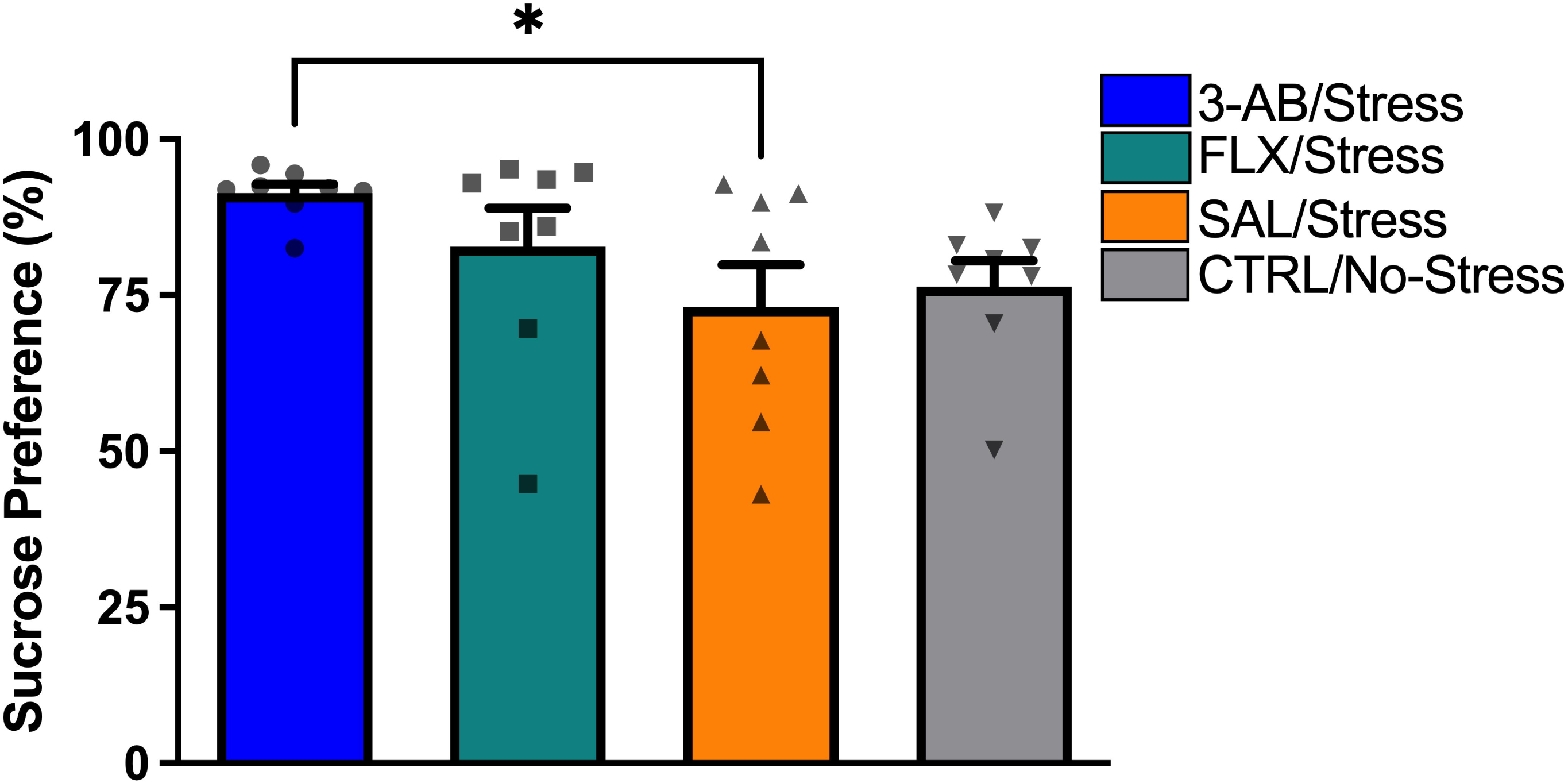
3-AB-treated stressed rats show higher sucrose preference than saline-treated rats. Sucrose preference (percent) measured on experimental days 8-10. The SAL/Stress group exhibited significantly lower preference compared to 3-AB/Stress (*p = 0.032) groups, indicating anhedonic behavior. Treatment with 3-AB prevented this stress-induced deficit. Individual data points represent each animal’s sucrose preference averaged across the 3-day testing period. Data are mean ± SEM (n = 8 per group).

### Social Interaction Test

Analysis of the social interaction test revealed a significant effect of Treatment on time spent in the avoidance zone [F(3,28) = 5.263, p = 0.005]. The SAL/Stress group (M = 65.41 ± 7.04%) spent significantly more time in the avoidance zone compared to the 3-AB/Stress group (M = 29.88 ± 12.16%, p < 0.045) and the CTRL/No-Stress group (M = 10.82 ± 3.38%, p < 0.01). The FLX/Stress group (M = 39.90 ± 13.56%) did not differ significantly from the SAL/Stress group (Fig 3).

**Figure 3.**
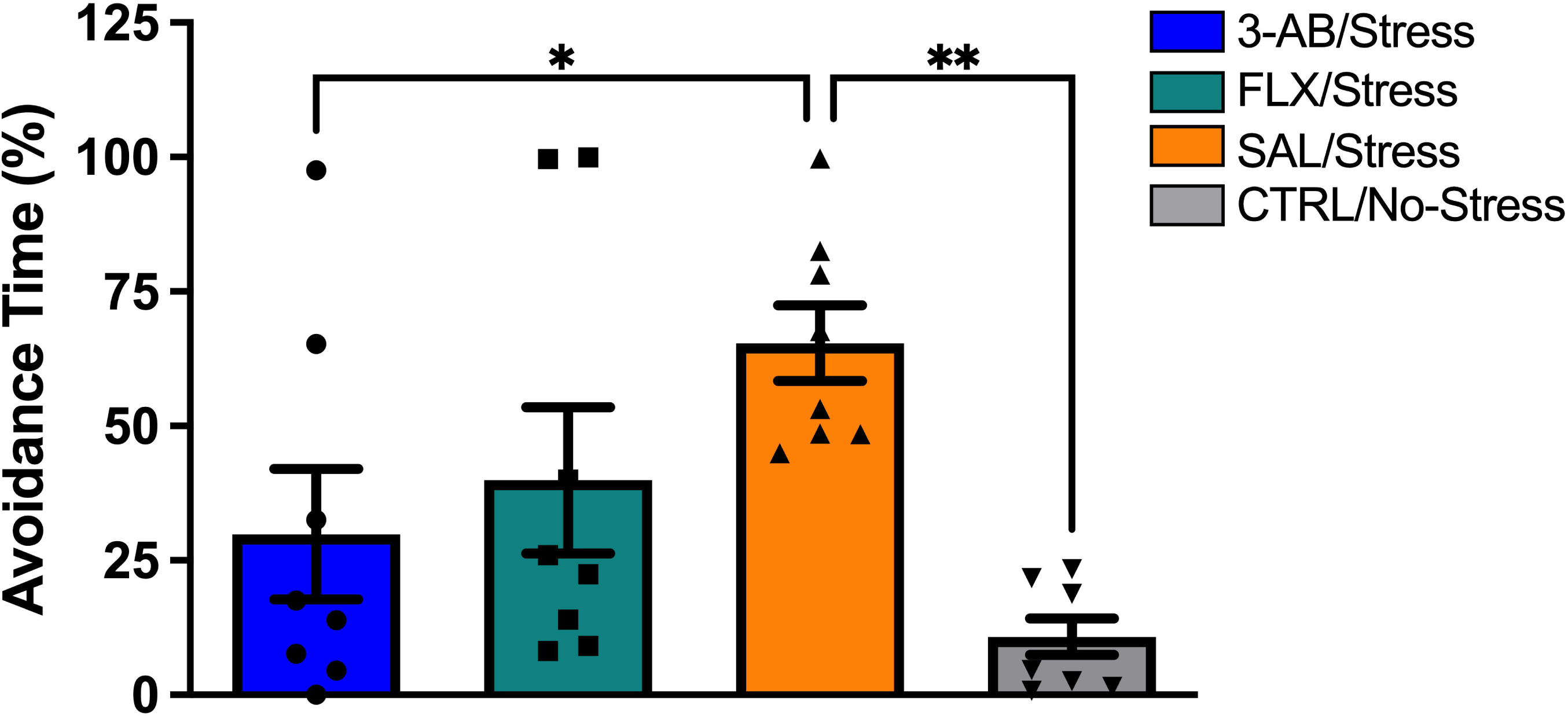
3-AB prevents stress-induced social avoidance. Time spent in the avoidance zone (percent) during the social interaction test on day 11. The SAL/Stress group exhibited significantly greater avoidance compared to 3-AB/Stress (*p < 0.05) and CTRL/No-Stress (**p < 0.01) groups. Treatment with 3-AB prevented this stress-induced social deficit. Individual data points represent individual animals. Data are mean ± SEM (n = 8 per group).

### Arterial Blood Pressure and Heart Rate Monitoring

To assess cardiovascular effects of chronic stress and potential protective effects of 3-AB, arterial blood pressure and heart rate were monitored before, during, and after the stress paradigm (Fig 4 & 5). Baseline hemodynamic parameters during the 3-day pre-stress period did not differ significantly between groups. Because treatment groups differed modestly at baseline for several measures, the Treatment x Time interaction (i.e., whether the change from baseline differed between groups) was used as the primary test of a treatment effect. (Fig 4).

**Figure 4.**
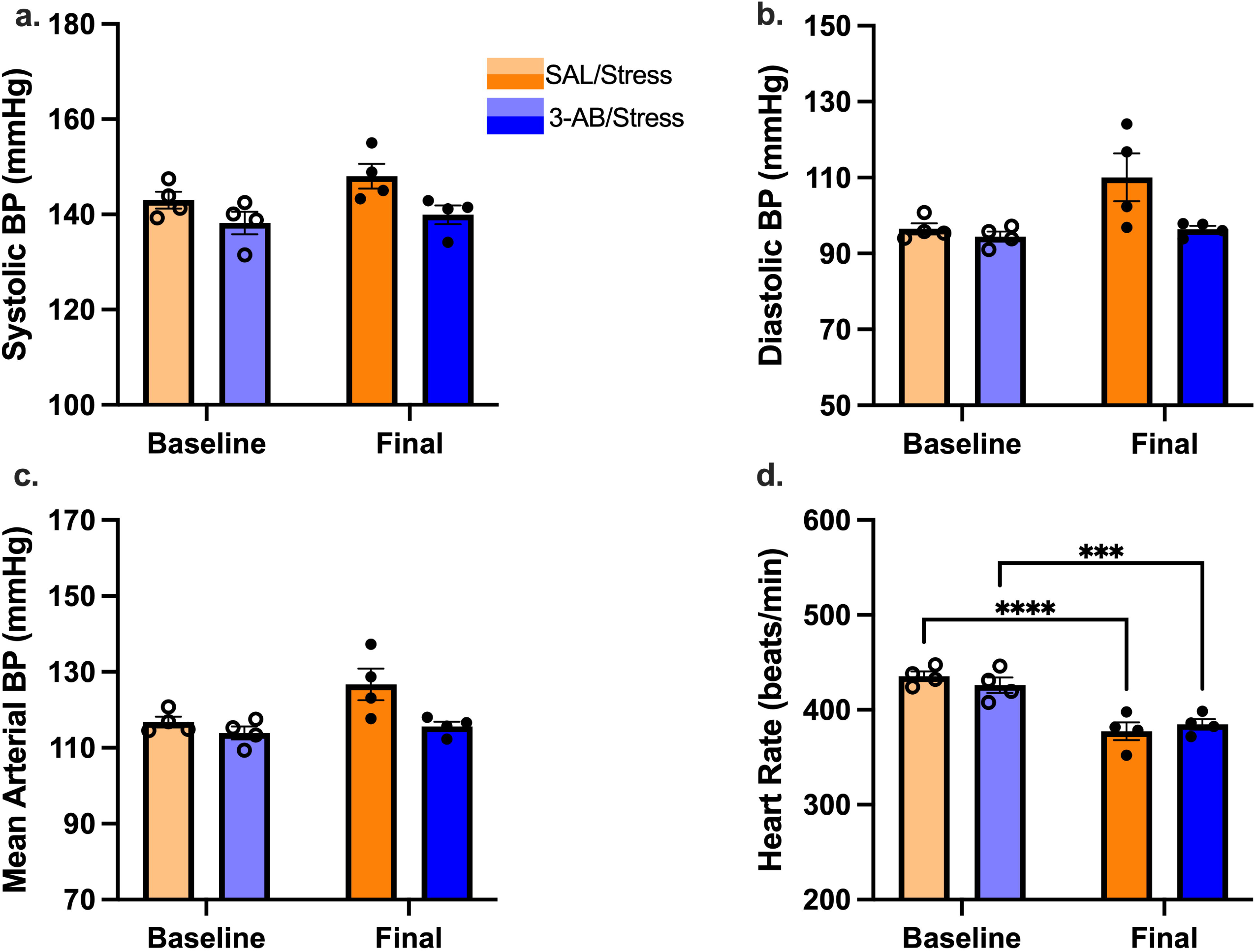
Hemodynamic changes across the stress paradigm in SAL/Stress and 3-AB Stress rats. Hemodynamic parameters measured via radiotelemetry before (baseline, open circles) and after (final, filled circles) the 10-day SDS+CUS paradigm. (A) Systolic blood pressure, (B) diastolic blood pressure, (C) mean arterial pressure, and (D) heart rate. Statistics are from two-way repeated-measures ANOVA; the Treatment x Time interaction tests whether the change from baseline differed between groups [SBP, DBP, and MAP interactions not significant; HR interaction p = 0.044 (*)]. Individual data points represent individual animal hemodynamic averages. Data are mean ± SEM (n = 4 per group).

### Systolic Blood Pressure

Two-way repeated measures ANOVA revealed a significant main effect of stress exposure (Time: baseline vs. post-stress) [F(1,6) = 9.68, p = 0.021], (Fig 4*a*), but no significant main effect of Treatment or Treatment × Time interaction effects (both p > 0.05). The estimated difference in change between SAL/Stress and 3-AB/Stress was 3.33 mmHG (95% CI -2.00 – 8.65), compatible with both no effect and a modest attenuation of the stress-induced rise. These data therefore do not establish that 3-AB altered the change in SBP.

### Diastolic Blood Pressure

Two-way repeated measures ANOVA revealed no significant main effects of Treatment or Time and no Treatment x Time interaction (Fig. 4*b*). The estimated difference in change between groups was 11.66 mmHg (95% CI -4.74 to 28.06). Variability in the SAL/Stress group was large (baseline: 96.5 ± 1.5 mmHg vs. post-stress: 110.1 ± 6.3 mmHG), reflecting marked increases in two of four rats, whereas 3-AB/Stress rats remained stable (94.5 ± 1.3 vs. 96.4 ± 0.9 mmHg).

### Mean Arterial Pressure

Analysis of mean arterial pressure showed a significant main effect of Treatment [F(1,6) = 6.91, p = 0.039] and Time [F(1,6) = 6.80, p = 0.040, Fig 4*c*] but no significant Treatment x Time interaction. The estimated difference in change between grops was 8.18 mmHg (95% CI -2.79 – 19.15). As with SBP, these data do not establish that 3-AB altered the change in MAP relative to SAL/Stress.

### Heart Rate

Heart rate declined in both groups (main effect of Time, F(1,6) = 233.4, p < 0.0001), with no main effect of Treatment and a significant interaction effect of Treatment × Time [F(1,6) = 6.484, p = 0.044, Figure 3d]. The decline was larger in SAL/Stress than in 3-AB/Stress rates (−58.0 ± 5.2 vs. -41.4 ± 3.9bpm; difference 16.6 bpm, 95% CI 0.7 to 32.5). Given the small sample and the number of hemodynamic measures tested, this finding should be considered exploratory.

### Acute Response to Social Defeat

Hemodynamics were recorded during social defeat. There were no significant differences between the SAL/Stress and 3-AB animals in systolic blood pressure, diastolic blood pressure, MAP, or heart rate (Fig 5*a-d*).

**Figure 5.**
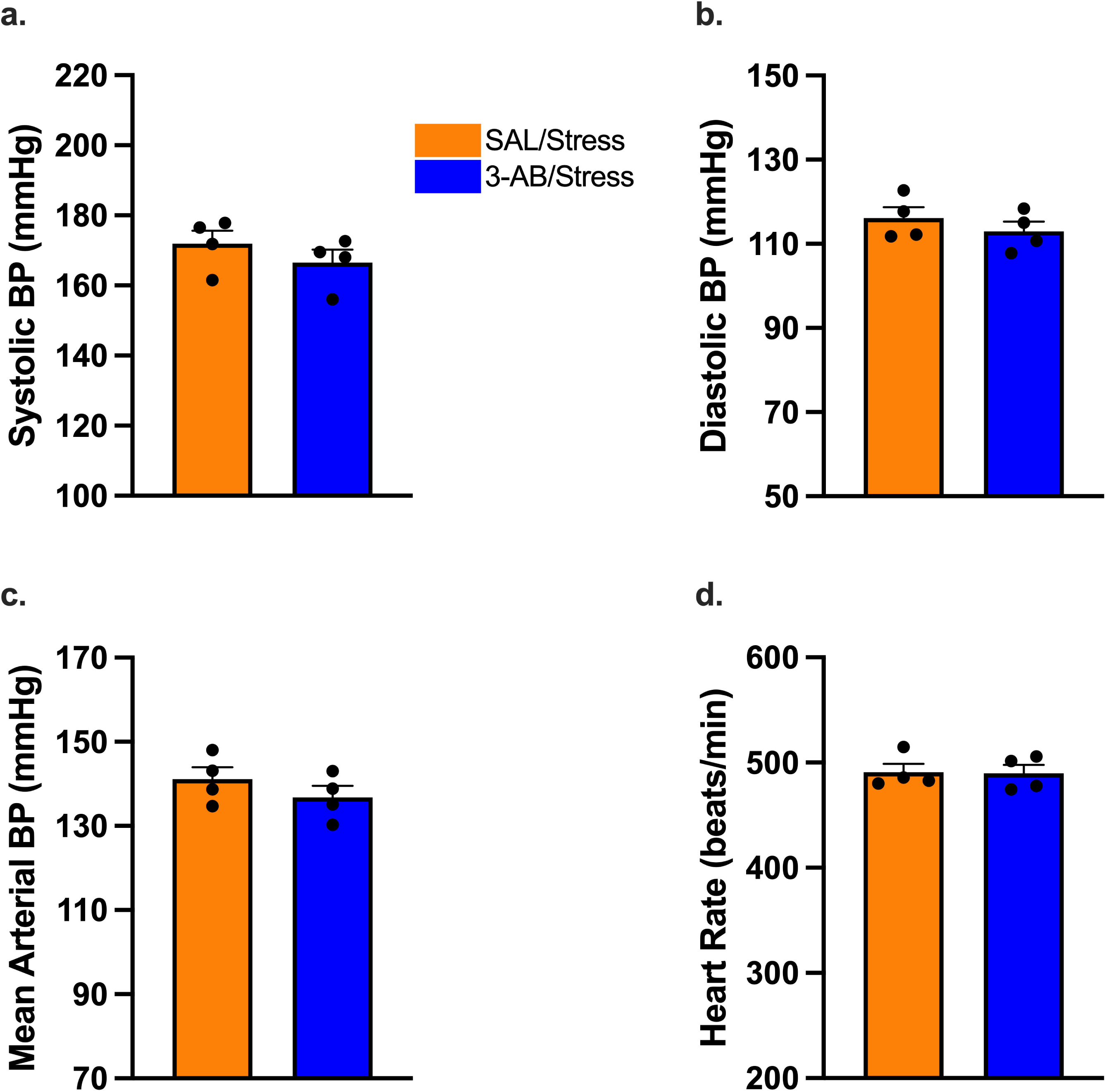
Acute hemodynamic responses during social defeat stress are similar between treatment groups. Real-time cardiovascular parameters measured via radiotelemetry during the 5-min social defeat stress sessions. (A) Systolic blood pressure, (B) diastolic blood pressure, (C) mean arterial pressure, and (D) heart rate during social defeat exposure. No significant differences were observed between SAL/Stress (orange) and 3-AB/Stress (blue) groups in any hemodynamic parameter during acute stress exposure. Individual data points represent individual animals. Data are mean ± SEM (n = 4 per group).

### PARP-1 Expression in Rat Brain Tissue

#### Prefrontal Cortical Gray Matter (L1 and L3-L6)

Since Brown-Forsythe tests indicated significant variance heterogeneity across groups in both L1 and L3-6, Welch’s ANOVA was used to assess group differences. No significant effect of Treatment was observed in L1 or L3-6 Fig 6*a-b*, respectively).

**Figure 6.**
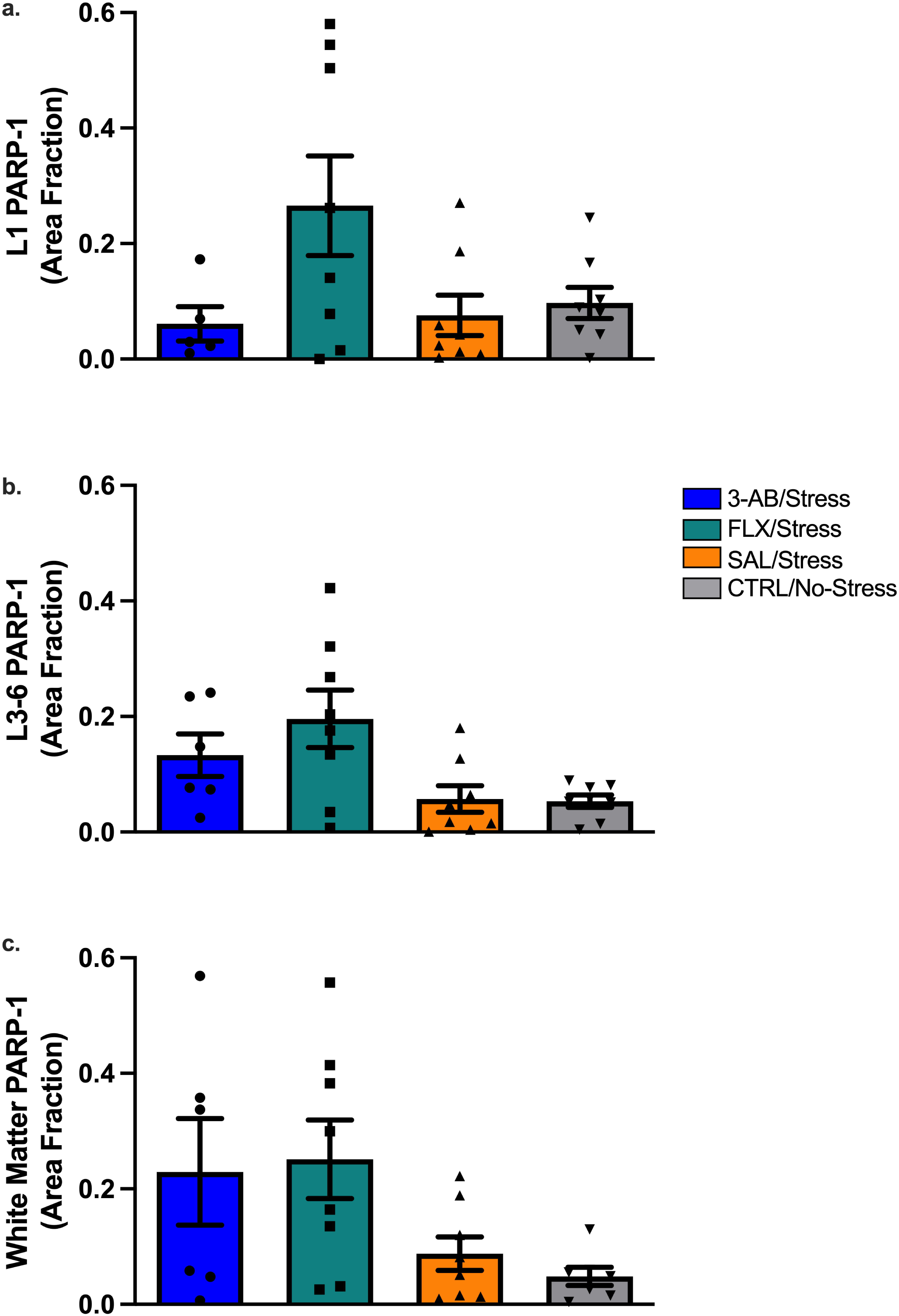
PARP-1 expression across treatment groups in prefrontal cortical gray and white matter. PARP-1 immunoreactivity (area fraction) in PFC subregions. (A) Layer 1 (L1) gray matter, (B) Layers 3-6 (L3-6) gray matter, and (C) forceps minor of the corpus callosum (white matter). Due to significant variance heterogeneity across groups (Brown-Forsythe test), Welch’s ANOVA was used for each region. No significant group differences were observed in L1 or L3-6. A significant main effect of Treatment was observed in FMI, but Dunnett’s T3 post-hoc comparisons showed no significant pairwise differences between groups when compared with SAL/Stress group. Individual data points represent individual animals. Data are mean ± SEM. Group sizes ranged from n = 5-8 across regions due to tissue availability (see Methods).

#### Forceps Minor of the Corpus Callosum

Brown-Forsythe testing also indicated significant variance heterogeneity in the FMI region; Welch’s ANOVA revealed a significant effect of Treatment [W(3, 11.8) = 3.69, p = 0.044]. However, Dunnett’s T3 post-hoc comparisons showed no significant differences between SAL/Stress group and the CTRL/No-Stress, FLX/Stress, or 3-AB/Stress groups (Fig 6*c*).

#### Microglial Morphology

IBA-1 immunohistochemistry (Fig. 7) and 3D morphological analysis of 15 microglia per animal (Fig 8, Supplementary Figs S1-3). Hierarchical linear mixed-effects models revealed significant effects of Treatment on microglial soma shape, specifically ellipticity (prolate) [F(3,28) = 3.227, p = 0.038; Fig 8*c*] and ellipticity (oblate) [F(3,28) = 4.507, p = 0.011; Fig. 8*d*]. Post-hoc comparisons using Holm-Šídák’s multiple comparisons test showed that microglia in the CTRL/No-Stress group were significantly less prolate (M = 0.400 ± 0.031) than those in the SAL/Stress group (M = 0.453 ± 0.036, p = 0.043), consistent with a surveilling rather than activated state (Fig 8*c*). Additionally, 3-AB/Stress group were significantly more oblate (M = 0.413 ± 0.026) than those in the SAL/Stress group (M = 0.367 ± 0.024, p = 0.007), indicating a less activated morphology (Fig 8*d*). Prolate and oblate ellipticity were significantly negatively correlated across animals (Person r = -0.614, 0.0002), consistent with these measures reflecting related dimensions of microglial soma shape change. No significant differences were observed in gray matter filament morphology or white matter microglial morphology between groups (Supplementary figs S1-3).

**Figure 7.**
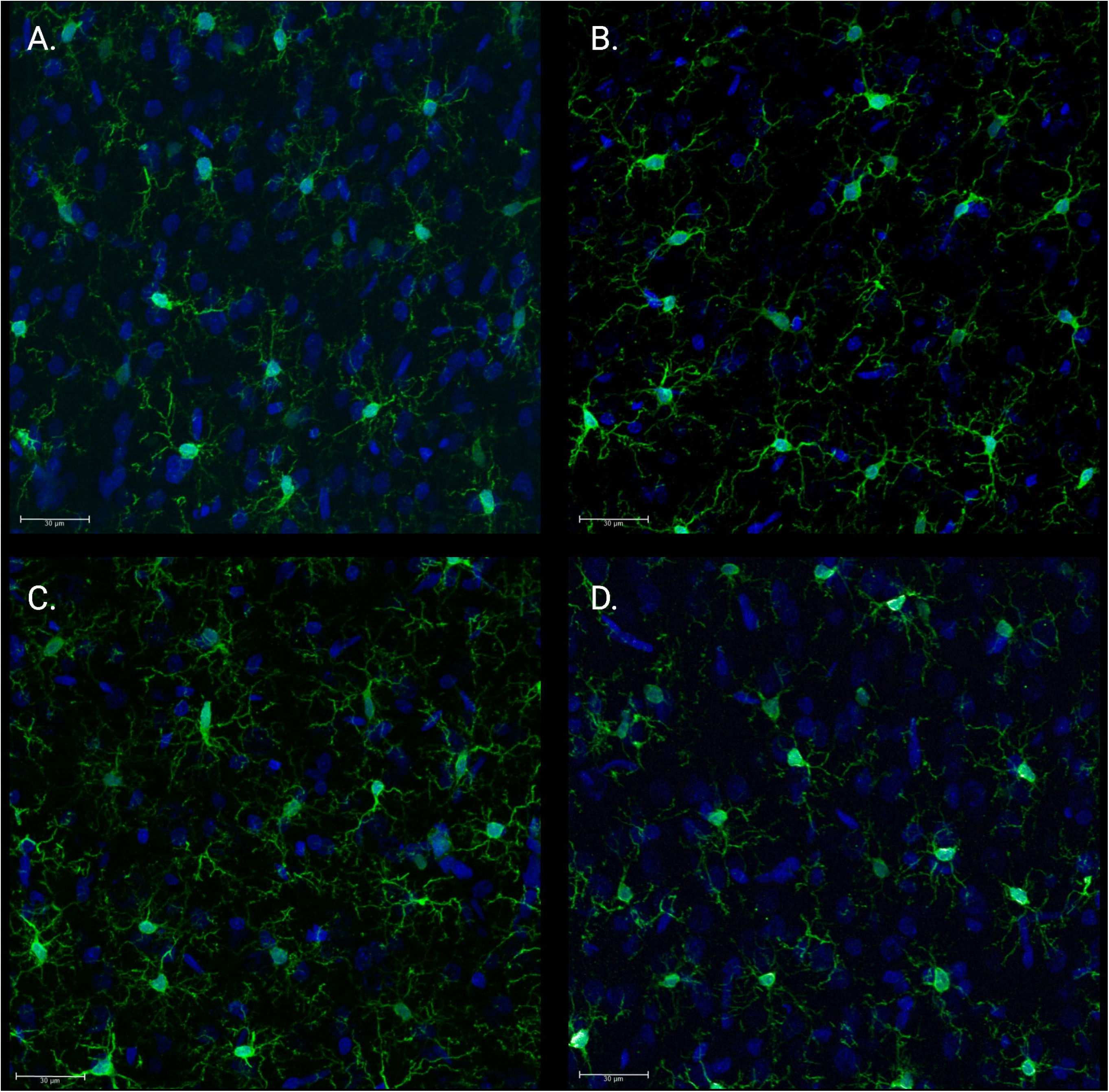
Microglial morphology in PFC varies by treatment. Microglia (green, Iba1) and nuclei (blue, DAPI) in PFC from (A) 3-AB/Stress, (B) FLX/Stress, (C) SAL/Stress, (D) CTRL/No-Stress groups. Morphological changes that can be observed across treatment groups, with SAL/Stress animals displaying more activated, prolate microglial phenotypes characterized by retracted processes and irregularly shaped soma compared to the ramified morphology observed in 3-AB/Stress, FLX/Stress, and CTRL/No-Stress groups. Scale bar = 30µm.

**Figure 8.**
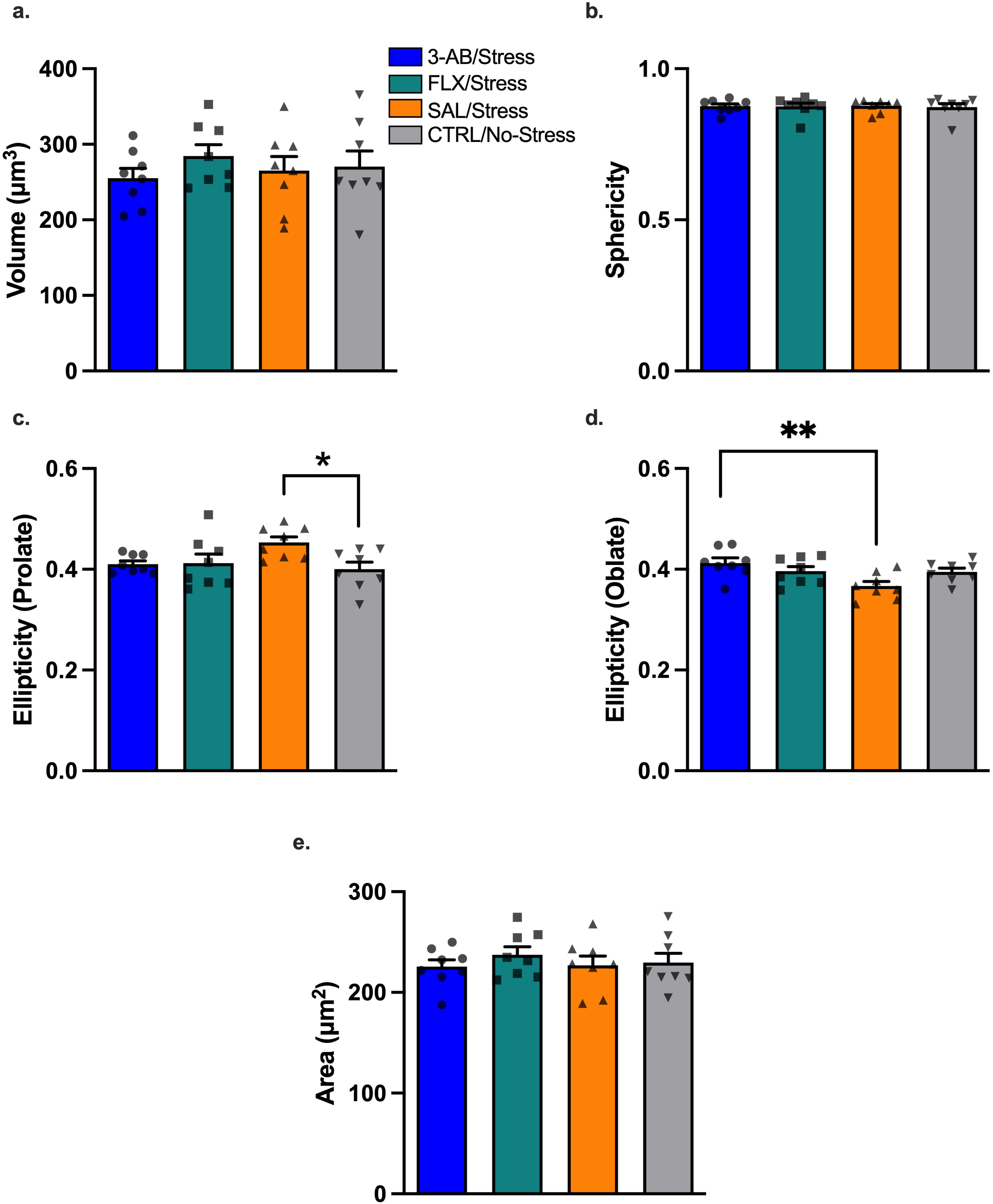
Microglial soma morphology in PFC gray matter. Microglial soma morphology parameters from 3D reconstruction of IBA-1-immunolabled microglia (15 cells per animal). (A) Soma volume, (B) soma sphericity, (C) soma ellipticity (prolate), (D) soma ellipticity (oblate), and (E) soma area. Chronic stress induced a more prolate (elongated) microglial soma shape in SAL/Stress rats compared to CTRL/No-Stress(*p = 0.043), consistent with microglial activation. Treatment with 3-AB prevented this stress-induced morphological change, with 3-AB/Stress microglia exhibiting significantly more oblate soma than SAL/Stress (**p = 0.007). Individual data points represent individual animals. Data are mean ± SEM (n = 8 per group).

#### Cytokine Expression

Analysis of proinflammatory cytokine levels in ventral HPC revealed significant effects of Treatment on TNF-α [F(3,28) = 5.35, p = 0.005], IL-1β [F(3,28) = 4.60, p < 0.010], and IL-6 [F(3,27) = 3.61, p = 0.026] expression (Fig 9). Brown-Forsythe testing confirmed homogeneity of variance for all three cytokines (all p’s > 0.16), supporting the use of ordinary ANOVA with Dunnett’s post-hoc comparisons.

**Figure 9.**
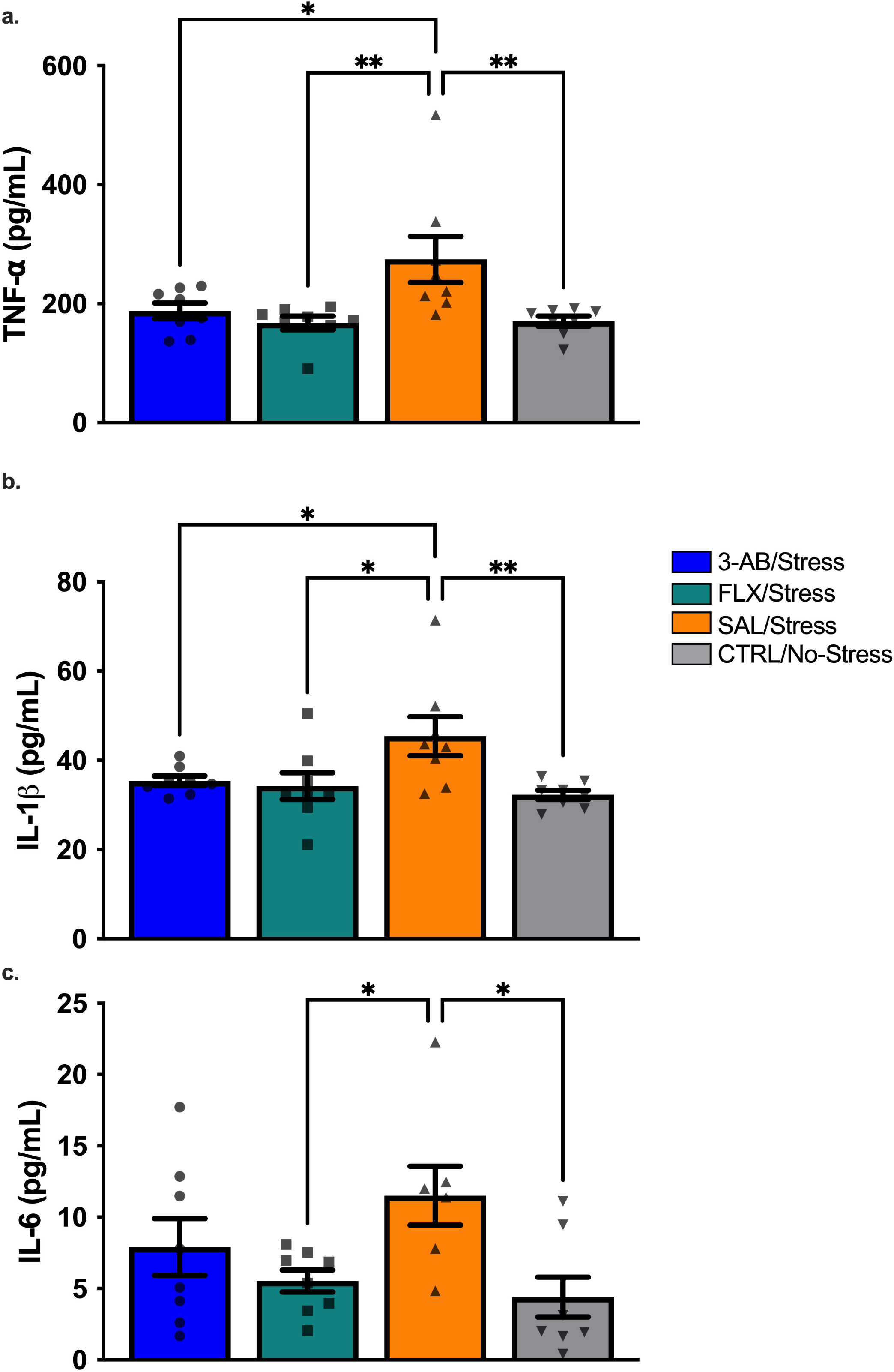
3-AB and FLX attenuate stress-induced increases in hippocampal proinflammatory cytokines. Hippocampal cytokine concentrations measured by ELISA. (A) Tumor necrosis factor-alpha (TNF-⍰), (B) interleukin-1 beta (IL-1β), and (C) interleukin-6 (IL-6). Chronic stress significantly elevated TNF-⍰ and IL-1β in SAL/Stress animals compared to 3-AB/Stress, FLX/Stress, and CTRL/No-Stress (*p < 0.05, **p < 0.01). For IL-6, Sal/Stress was significantly elevated compared to CTRL/No-Stress and FLX/Stress animals (*p < 0.05). One SAL/Stress sample was unavailable for IL-6 analysis (n = 7); all other groups/ analytes, n = 8 per group. Individual data points represent individual animals. Data are mean ± SEM.

#### TNF-α

The SAL/Stress group exhibited significantly higher TNF-α levels compared to the 3-AB/Stress (mean difference = 86.72 pg/mL, p = 0.025), FLX/Stress (mean difference = 106.8 pg/mL, 95%, p = 0.005), and CTRL/No-Stress (mean difference = 105.8 pg/mL, p = 0.006) groups (Fig 9*a*).

#### IL-1β

The SAL/Stress group exhibited significantly higher IL-1β levels compared to the 3-AB/Stress (mean difference = 10.01 pg/mL, p = 0.040), FLX/Stress (mean difference = 11.15pg/mL, p = 0.020), and CTRL/No-Stress (mean difference = 13.20 pg/mL, p = 0.006) groups (Fig 9*b*).

#### IL-6

The SAL/Stress group demonstrated significantly higher IL-6 levels compared to the FLX/Stress (mean difference = 5.97 pg/mL, p = 0.041) and CTRL/No-Stress (mean difference = 7.09 pg/mL, p = 0.020) groups but did not significantly differ from the 3-AB/Stress group (mean difference = 3.59 pg/mL) (Fig 9*c*).

## Discussion

The present study investigated the effects of the PARP-1 inhibitor 3-AB in a rodent model of chronic psychological stress while proposing possible neurobiological mechanisms underlying these effects. Our findings suggest that 3-AB produces antidepressant-like behavioral effects while attenuating select neuroinflammatory responses to chronic stress. The precise molecular mechanisms remain to be established. The absence of significant PARP-1 upregulation in SAL/Stress group relative to controls was specific to the PFC and timepoint assessed here and does not exclude PARP-1 involvement in other brain regions (e.g., HPC) or at other post-stress intervals. Our reasoning for testing the PFC was due to our previous findings in human postmortem tissue demonstrating increase PARP-1 expression in MDD tissue compared to healthy controls^20^. Given this limitation, we cannot directly attribute the effects of 3-AB to PARP-1 inhibition in this study, and alternative or additional mechanisms should be considered in future work.

### Behavioral Effects of 3-AB

The sucrose preference test, a well-established measure of anhedonia in rodents^34^, revealed that, averaged across the testing period, 3-AB-treated stressed rats showed significantly higher sucrose preference than saline-treated rats. In contrast, sucrose preference in CTRL/No-Stress and FLX/Stress groups did not differ significantly from Sal/Stress group. This unexpected outcome in the CTRL/No-Stress group may reflect acute isolation stress, as CTRL animals were group-housed through the study but required brief single-housing during testing. That transition is known to induce stress and potentially confound baseline sucrose consumption^35^. Similarly, in the social interaction test, 3-AB treatment prevented the development of social avoidance behavior in stressed rats. These findings corroborate previous work from our laboratory demonstrating antidepressant-like effects of 3-AB in rodent models^23^ .

Interestingly, while 3-AB consistently prevented stress-induced behavioral deficits, FLX treatment failed to produce significant improvements in either sucrose preference or social interaction. For sucrose preference, this finding is contrary to our previous work^23^, in which FLX significantly improved this measure. This behavioral inconsistency is less unexpected for social interaction, however, as FLX’s effect on this measure in our prior work ^23^ also did not reach significance. As this protocol closely parallels our prior work, in which fluoxetine was at least partially effective, we consider this discordant result more likely to reflect procedural or paradigm-specific factors than confirmed treatment resistance^4,36^. This differential response highlights the potential value of further characterizing 3-AB’s mechanism of action relative to established antidepressants.

### Chronic Stress and Hemodynamic Changes

Chronic stress was associated with increases in SBP and MAP across the paradigm, and DBP rose in some saline-treated rats. These increases tended to be smaller in 3-AB-treated rats but the Treatment x Time interactions were not significant and the confidence intervals for the between-groups differences in change include zero (e.g., SBP: 3.33 mmHG, 95% CI -2.00 – 8.65). These data therefore do not provide statistical evidence that 30AB altered the blood pressure response to chronic stress, although with four animals per group the study had limited power to detect a modest effect. These observations are consistent with prior work linking chronic stress to cardiovascular dysfunction ^15–17^ and they motivate future investigation with larger cohorts. Heart rate declined in both groups, with a greater decline in SAL/Stress group; this was the only hemodynamic measure with a nominally significant interaction and should be interpreted cautiously. One possible contributor to the cardiovascular protective effects of 3-AB is preservation of autonomic nervous system function, although this was not directly assessed in the present study. In a distinct model of dyslipidemia-induced vascular dysfunction, PARP-1 deletion has been shown to restore baroreflex sensitivity via reduced oxidative stress and inducible nitric oxide (iNOS) upregulation ^37^, and PARP-1 overactivation has more broadly been implicated in endothelial dysfunction and vascular inflammation^38,39^. Whether comparable oxidative or autonomic mechanisms contribute to the cardiovascular effects observed here is unknown. Finally, both groups showed similar acute hemodynamic responses during social defeat sessions (Fig. 5), indicating that 3-AB did not impair normal autonomic reactivity to acute stressors.

### PARP-1 Expression and Fluoxetine Interaction

Contrary to our original hypothesis, we did not find statistically robust group differences in PARP-1 expression in PFC gray or white matter. Crucially, however, a lack of change in baseline PARP-1 protein abundance does not preclude its functional enzymatic involvement in SDS+CUS. PARP-1 is a constitutive enzyme with low baseline activity, but it can undergo massive catalytic hyperactivation in response to cellular stress. When triggered by DNA damage or metabolic shifts, its enzymatic rate can increase over 100-fold to drive downstream poly (ADP-ribose) polymer synthesis^40^. As 3-AB acts as a competitive inhibitor of catalytic activity rather than a suppressor of protein translation^41^, it is possible that a stress-induced increase PARP-1 enzymatic activity occurred without a corresponding change in protein expression detectable by IHC. Future studies will further investigate this possibility. Notably, however, PARP-1 expression variance differed systematically by treatment; within each region examined, variability was consistently lowest in the CTRL/No-Stress group, intermediate in the SAL/Stress group, and highest in the FLX/Stress group (Fig. 6). This pattern was not observed for 3-AB/Stress, which showed variance comparable to or lower than SAL/Stress in cortical gray matter, though more variable in white matter.

This FLX-specific increase in inter-individual variability may itself be an informative finding rather than a limitation of measurement. It suggests that FLX treatment during chronic stress introduces individual variability in PARP-1 regulation that is not observed with stress exposure alone or with 3-AB treatment. This is consistent with our behavioral findings, in which FLX/Stress animals showed heterogeneous, non-significant responses on both sucrose preference and social interaction measures, in contrast to the more uniform effects observed in the 3-AB/Stress group. Together, these findings suggests that FLX’s effects on PARP-1 expression, like its behavioral effects, may be more variable across individual animals than those of 3-AB. This individual variability could help explain why FLX did not produce consistent behavioral efficacy in this paradigm.

The absence of significant group differences in PARP-1 protein expression may also reflect methodological factors. Our combined 10-day SDS+CUS model has previously been shown to significantly elevate DNA oxidation in PFC white matter, but not gray matter, in stressed rats relative to controls^20^, indicating that this paradigm reliably produces upstream oxidative damage even when downstream PARP-1 protein expression is not detectably altered. Furthermore, elevated PARP1 gene expression in MDD has previously been detected in specific glial cell populations using laser-capture microdissection combined with quantitative PCR ^20^, a more sensitive and cell-type-specific approach than the whole-tissue IHC area fraction method used here. Given that PARP1 activity is also subject to rapid turnover via poly(ADP-ribose) glycohydrolase-mediated degradation, it is plausible that transient or cell-type-specific changes in PARP-1 were not captured by our IHC approach at the single 48h timepoint examined. Future studies using direct activity assays (such as PAR accumulation ELISAs) with larger sample sizes will be needed to determine whether this individual variability in PARP-1 expression relates to differential treatment response and whether it reflects a broader pattern of heterogeneous FLX response under conditions of chronic stress.

### Microglial Activation and Neuroinflammation

Our analysis of microglial morphology revealed that stressed rats treated with saline exhibited significantly more prolate (elongated) microglia in PFC gray matter compared to non-stressed controls, consistent with a shift along the morphological spectrum associated with microglial activation, though morphology alone does not confirm activation state^42^. In contrast, stressed rats treated with 3-AB displayed significantly more oblate microglia, resembling the morphology seen in control rats^42^ . These findings suggest that 3-AB may exert its antidepressant-like effects in part by preventing this stress-induced microglial morphological shift, though morphology alone cannot confirm the functional activation state of these cells.

The lack of significant morphological changes in white matter microglia was unexpected, given our previous finding of increased oxidative DNA damage in PFC white matter following chronic stress^20,23^ . This discrepancy may be due to the timing of tissue collection (48 h after the final stressor), allowing microglia to return to a surveilling state. Future studies should examine microglial activation at various time points following stress exposure to capture the temporal dynamics of microglial responses to chronic stress.

### Proinflammatory Cytokine Expression

Analysis of proinflammatory cytokines in the ventral HPC revealed that 3-AB treatment significantly attenuated the stress-induced increases in both TNF-α and IL-1β, with no significant effect on IL-6. These findings are consistent with previous research demonstrating anti-inflammatory effects of PARP inhibitors^10^ and suggest a possible contributing mechanism for 3-AB’s antidepressant-like effects.

Proinflammatory cytokines have been implicated in the pathophysiology of depression through various mechanisms, including disruption of monoamine neurotransmission^43^, activation of the kynurenine pathway^44^, and promotion of oxidative stress^11^. By attenuating stress-induced increase in TNF-α and IL-1β, 3-AB may prevent some of these detrimental effects on neural function and mood regulation, although the lack of effect on IL-6 limits conclusions about broad anti-inflammatory action.

### Proposed Mechanism and Clinical Implications

Based on our findings, we hypothesize that 3-AB may exert antidepressant-like effects at least in part through inhibition of PARP-1-mediated neuroinflammatory processes, though this study did not directly demonstrate PARP-1 inhibition. One possibility is that by limiting PARP-1 activation, 3-AB could reduce NF-κB-dependent transcription of proinflammatory genes, thereby decreasing the microglial morphological shift, TNF-α, and IL-1β production observed. The prevention of stress-induced hypertension by 3-AB treatment indicates that it provides multi-system benefits that may be particularly valuable in treating the complex pathophysiology of stress-related disorders and increased risk of cardiovascular disease in this population. Exploratory analyses of gut microbiome composition revealed stress-induced alterations that were partially prevented by 3-AB treatment, further supporting multi-system effects (See Supplemental Materials).

These preclinical findings may have clinical relevance worth further exploration. PARP inhibitors are already approved for cancer treatment, which could facilitate future investigation of repurposing for psychiatric indications. A case report from our laboratory describing rapid antidepressant and anxiolytic effects following niraparib administration in a patient with treatment-resistant depression^24^ is an intriguing observation that warrants further investigation in the clinical population.

### Exploratory Microbiome Findings

Although not the primary focus of this study, exploratory microbiome analyses were conducted to assess potential peripheral effects of chronic stress and PARP inhibition (see Supplemental Materials). Overall, microbial community composition (beta-diversity) did not differ significantly among groups. However, a selective reduction in *Bacteroides dorei* abundance was observed in SAL/Stress rats compared with both the CTRL/No-Stress group and the 3-AB/Stress group, while no difference was observed between the CTRL/No-Stress and 3-AB/ Stress groups (Supplementary Table 1). This finding suggests that chronic stress selectively affects *B. dorei* and that 3-AB was associated with the preservation of levels comparable to those in controls. *Bacteroides* species have been implicated in promoting gut health through protective measures, including modulation of detoxifying peroxidases as part of the antioxidant response^45^. These pathways are particularly relevant given the antioxidant deficits we have demonstrated in human stress-related disorders^20,46,47^.

An exploratory analysis of microbial metabolites identified reduced valeric acid levels in SAL/Stress animals compared with CTRL/No-Stress animals (Supplemental Fig. 4f), while 3-AB-treated animals showed intermediate levels that did not differ significantly from either group. Previous studies have reported reduced valeric acid following chronic stress and have shown beneficial effects of short-chain fatty acids on stress-behavioral outcomes ^48,49^.

These exploratory findings do not demonstrate broad alterations of the gut microbiota but instead suggest that chronic stress selectively alters specific microbial features and that 3-AB may prevent some of the stress-associated changes. Future studies specifically designed and adequately powered for comprehensive microbiome analyses will be required to determine whether PARP inhibition exerts a broader effect.

### Future Directions

Several important questions remain to be addressed in future research. First, our study focused exclusively on male rats, despite the higher prevalence of MDD in women. Sex differences in stress responses and neuroinflammatory processes are well-documented^50,51^, and future studies should include female rats to assess potential sex-specific effects of PARP inhibition. Second, this study did not include measurement of corticosterone or serotonin, which would provide additional mechanistic insight into hypothalamic-pituitary-adrenal axis activity and monoaminergic contributions to the observed phenotypes; these measures are planned for future work. Third, the timing of tissue collection (48h after the final stressor) may have allowed for partial resolution of neuroinflammatory responses, potentially obscuring some acute effects. Time-course studies examining neuroinflammatory markers at multiple time points following stress exposure would provide valuable insights into the temporal dynamics of stress-induced pathophysiology and PARP inhibitor effects. Fourth, future research should investigate the long-term effects of PARP inhibitor treatment, potential interactions with traditional antidepressants, and effects on other neurobiological systems implicated in depression, such as neurotrophic signaling and neurogenesis. Additionally, the marked inter-individual variability in PARP-1 expression observed specifically in the FLX/Stress group warrants further investigation, including larger sample sizes and direct measures of PARP-1 enzymatic activity, to determine whether this reflects heterogeneous treatment response to fluoxetine under conditions of chronic stress.

## Conclusion

This study provides evidence for the antidepressant-like effects of 3-AB in a rodent model of chronic psychological stress. Our findings demonstrate that 3-AB treatment prevented stress-induced behavioral deficits and cardiovascular dysfunction while selectively attenuating stress-induced changes in microglial morphology and hippocampal TNF-α and IL-1β. These results are consistent with a role for neuroinflammatory processes in stress-related pathophysiology, though our data does not establish PARP-1 inhibition as the specific mechanism underlying these effects. Future studies incorporating direct measures of PARP-1 enzymatic activity, additional behavioral assessments, and female subjects will be needed to clarify the mechanism of action of 3-AB and its potential relevance to stress-related psychiatric disorders.

## Note

Supplemental materials include microglial morphology graphs for gray matter filaments, white matter soma and filaments, exploratory analyses of gut microbiome composition (16S rRNA sequencing), short-chain fatty acid quantification, with detailed methods and results available online.

## Acknowledgments

None.

## Data Availability Statement

The data that support the findings of this study are available from the corresponding author upon reasonable request.

## Ethical Statement

All animal procedures were conducted in accordance with the NIH Guide for the Care and Use of Laboratory Animals and were approved by the East Tennessee State University Institutional Animal Care and Use Committee.

## References

1. König, H., König, H.-H. & Konnopka, A. The excess costs of depression: a systematic review and meta-analysis. Epidemiology Psychiatr. Sci. 29, e30 (2019).

2. Roman, M. & Irwin, M. R. Novel neuroimmunologic therapeutics in depression: A clinical perspective on what we know so far. *Brain, Behav.*, Immun. 83, 7–21 (2020).

3. Akil, H. et al. Treatment resistant depression: A multi-scale, systems biology approach. Neurosci. Biobehav. Rev. 84, 272–288 (2018).

4. Rush, A. J. The varied clinical presentations of major depressive disorder. J. Clin. psychiatry 68 **Suppl 8**, 4–10 (2007).

5. Irwin, M. R. & Miller, A. H. Depressive disorders and immunity: 20 years of progress and discovery. *Brain, Behav.*, Immun. 21, 374–383 (2007).

6. Swardfager, W., Rosenblat, J. D., Benlamri, M. & McIntyre, R. S. Mapping inflammation onto mood: Inflammatory mediators of anhedonia. Neurosci. Biobehav. Rev. 64, 148–166 (2016).

7. Raison, C. L., Capuron, L. & Miller, A. H. Cytokines sing the blues: inflammation and the pathogenesis of depression. Trends Immunol. 27, 24–31 (2006).

8. Dantzer, R. Cytokine, Sickness Behavior, and Depression. Immunol. Allergy Clin. North Am. 29, 247–264 (2009).

9. Libby, P., Mallat, Z. & Weyand, C. Immune and inflammatory mechanisms mediate cardiovascular diseases from head to toe. Cardiovasc. Res. 117, 2503–2505 (2021).

10. Sriram, C. S. et al. Poly (ADP-ribose) polymerase-1 inhibitor, 3-aminobenzamide pretreatment ameliorates lipopolysaccharide-induced neurobehavioral and neurochemical anomalies in mice. Pharmacol. Biochem. Behav. 133, 83–91 (2015).

11. Herbet, M. et al. Altered expression of genes involved in brain energy metabolism as adaptive responses in rats exposed to chronic variable stress; changes in cortical level of glucogenic and neuroactive amino acids. Mol. Med. Rep. 19, 2386–2396 (2019).

12. Frank, M. G., Fonken, L. K., Watkins, L. R. & Maier, S. F. Microglia: Neuroimmune-sensors of stress. Semin. Cell Dev. Biol. 94, 176–185 (2019).

13. Nelson, L. H., Saulsbery, A. I. & Lenz, K. M. Small cells with big implications: Microglia and sex differences in brain development, plasticity and behavioral health. Prog Neurobiol 176, 103–119 (2019).

14. Rahimian, R., Belliveau, C., Chen, R. & Mechawar, N. Microglial Inflammatory-Metabolic Pathways and Their Potential Therapeutic Implication in Major Depressive Disorder. Front. Psychiatry 13, 871997 (2022).

15. Kim, H.-G., Cheon, E.-J., Bai, D.-S., Lee, Y. H. & Koo, B.-H. Stress and Heart Rate Variability: A Meta-Analysis and Review of the Literature. Psychiatry Investig. 15, 235–245 (2018).

16. Ajibewa, T. A. et al. Chronic Stress and Cardiovascular Events: Findings From the CARDIA Study. Am. J. Prev. Med. 67, 24–31 (2024).

17. Steptoe, A. & Kivimäki, M. Stress and cardiovascular disease. Nat. Rev. Cardiol. 9, 360– 370 (2012).

18. Bai, P. & Virág, L. Role of poly(ADP ribose) polymerases in the regulation of inflammatory processes. FEBS Lett. 586, 3771–3777 (2012).

19. Hassa, P. O. & Hottiger, M. O. A Role of Poly (ADP-Ribose) Polymerase in NF-B Transcriptional Activation. Biol. Chem. 380, 953–959 (1999).

20. Szebeni, A. et al. Elevated DNA Oxidation and DNA Repair Enzyme Expression in Brain White Matter in Major Depressive Disorder. Int J Neuropsychoph 20, pyw114 (2016).

21. Ha, H. C., Hester, L. D. & Snyder, S. H. Poly(ADP-ribose) polymerase-1 dependence of stress-induced transcription factors and associated gene expression in glia. Proc. Natl. Acad. Sci. 99, 3270–3275 (2002).

22. Kauppinen, T. M. & Swanson, R. A. Poly(ADP-Ribose) Polymerase-1 Promotes Microglial Activation, Proliferation, and Matrix Metalloproteinase-9-Mediated Neuron Death. J. Immunol. 174, 2288–2296 (2005).

23. Ordway, G. A. et al. Antidepressant-Like Actions of Inhibitors of Poly(ADP-Ribose) Polymerase in Rodent Models. Int J Neuropsychoph 20, 994–1004 (2017).

24. Jewett, B. E. et al. Rapid and temporary improvement of depression and anxiety observed following niraparib administration: a case report. BMC Psychiatry 20, 171 (2020).

25. Munshi, S. et al. Repeated stress induces a pro-inflammatory state, increases amygdala neuronal and microglial activation, and causes anxiety in adult male rats. *Brain, Behav.*, Immun. 84, 180–199 (2020).

26. Pryce, C. R. & Fuchs, E. Chronic psychosocial stressors in adulthood: Studies in mice, rats and tree shrews. Neurobiol. Stress 6, 94–103 (2017).

27. Rygula, R., Abumaria, N., Domenici, E., Hiemke, C. & Fuchs, E. Effects of fluoxetine on behavioral deficits evoked by chronic social stress in rats. Behav. Brain Res. 174, 188–192 (2006).

28. Flannelly, K. & Lore, R. The influence of females upon aggression in domesticated male rats (Rattus norvegicus). Anim. Behav. 25, 654–659 (1977).

29. Bondi, C. O., Rodriguez, G., Gould, G. G., Frazer, A. & Morilak, D. A. Chronic Unpredictable Stress Induces a Cognitive Deficit and Anxiety-Like Behavior in Rats that is Prevented by Chronic Antidepressant Drug Treatment. Neuropsychopharmacology 33, 320– 331 (2008).

30. Willner, P., Towell, A., Sampson, D., Sophokleous, S. & Muscat, R. Reduction of sucrose preference by chronic unpredictable mild stress, and its restoration by a tricyclic antidepressant. Psychopharmacology 93, 358–364 (1987).

31. Accardi, M. V. et al. Rat cardiovascular telemetry: Marginal distribution applied to positive control drugs. J. Pharmacol. Toxicol. Methods 81, 120–127 (2016).

32. Kaur, S. et al. In Vivo Telemetry to Record Long-Term Cardiovascular Parameters, Temperature, and Activity in Spinal Cord Injury Rat Models. J. Vis. Exp. : JoVE 10.3791/69714 (2026) doi:10.3791/69714.

33. D’Aquila, PaoloS., Newton, J. & Willner, P. Diurnal Variation in the Effect of Chronic Mild Stress on Sucrose Intake and Preference. Physiol. Behav. 62, 421–426 (1997).

34. Liu, M.-Y. et al. Sucrose preference test for measurement of stress-induced anhedonia in mice. Nat. Protoc. 13, 1686–1698 (2018).

35. Primo, M. J., Fonseca-Rodrigues, D., Almeida, A., Teixeira, P. M. & Pinto-Ribeiro, F. Sucrose preference test: A systematic review of protocols for the assessment of anhedonia in rodents. Eur. Neuropsychopharmacol. 77, 80–92 (2023).

36. Williams, L. M., Debattista, C., Duchemin, A.-M., Schatzberg, A. F. & Nemeroff, C. B. Childhood trauma predicts antidepressant response in adults with major depression: data from the randomized international study to predict optimized treatment for depression. Transl. Psychiatry 6, e799–e799 (2016).

37. Hans, C. P. et al. Protective Effects of PARP-1 Knockout on Dyslipidemia-Induced Autonomic and Vascular Dysfunction in ApoE−/− Mice: Effects on eNOS and Oxidative Stress. PLoS ONE 4, e7430 (2009).

38. Szabó, C. et al. Poly(ADP-Ribose) Polymerase Is Activated in Subjects at Risk of Developing Type 2 Diabetes and Is Associated With Impaired Vascular Reactivity. Circulation 106, 2680–2686 (2002).

39. Pacher, P. et al. The Role of Poly(ADP-Ribose) Polymerase Activation in the Development of Myocardial and Endothelial Dysfunction in Diabetes. Diabetes 51, 514–521 (2002).

40. Pandey, N. & Black, B. E. Rapid Detection and Signaling of DNA Damage by PARP-1. Trends Biochem. Sci. 46, 744–757 (2021).

41. Shieh, W. M. et al. Poly(ADP-ribose) Polymerase Null Mouse Cells Synthesize ADP-ribose Polymers*. J. Biol. Chem. 273, 30069–30072 (1998).

42. Kettenmann, H., Hanisch, U.-K., Noda, M. & Verkhratsky, A. Physiology of Microglia. Physiol. Rev. 91, 461–553 (2011).

43. Miller, A. H. & Raison, C. L. The role of inflammation in depression: from evolutionary imperative to modern treatment target. Nat. Rev. Immunol. 16, 22–34 (2016).

44. Réus, G. Z. et al. The role of inflammation and microglial activation in the pathophysiology of psychiatric disorders. Neuroscience 300, 141–154 (2015).

45. Zafar, H. & Saier, M. H. Gut Bacteroides species in health and disease. Gut Microbes 13, 1848158 (2021).

46. Chandley, M. J. et al. Markers of elevated oxidative stress in oligodendrocytes captured from the brainstem and occipital cortex in major depressive disorder and suicide. Prog. Neuro-Psychopharmacol. Biol. Psychiatry 117, 110559 (2022).

47. Szebeni, A. et al. Shortened telomere length in white matter oligodendrocytes in major depression: potential role of oxidative stress. Int. J. Neuropsychopharmacol. 17, 1579–1589 (2014).

48. Chen, M., Wang, L., Lou, Y. & Huang, Z. Effects of chronic unpredictable mild stress on gut microbiota and fecal amino acid and short-chain fatty acid pathways in mice. Behav. Brain Res. 464, 114930 (2024).

49. Wouw, M. van de et al. Short chain fatty acids: microbial metabolites that alleviate stress induced brain–gut axis alterations. J. Physiol. 596, 4923–4944 (2018).

50. Maeng, L. Y. & Milad, M. R. Sex differences in anxiety disorders: Interactions between fear, stress, and gonadal hormones. Horm. Behav. 76, 106–117 (2015).

51. Rainville, J. R. & Hodes, G. E. Inflaming sex differences in mood disorders. Neuropsychopharmacology 44, 184–199 (2019).

