## Supplementary Materials for "Behavioral, Cardiovascular, and Neuroinflammatory Effects of 3-Aminobenzamide in a Rodent Model of Chronic Psychological Stress"

**Supplementary Material**

**Supplemental Methods for Microbiome Analysis**

***DNA purification and microbiome sequencing***

Feces were collected from rats (n=8 per CTRL/No Stress, CTRL/Stress and 3-AB/Stress groups) immediately prior to euthanasia in clean, dry cages. Additionally, luminal samples were harvested from the colon post-euthanasia. Samples were stored at -80^o^C until processing. Genomic DNA was extracted from approximately 100 mg of fecal or colonic tissue using the DNeasy PowerSoil Pro Kit (Qiagen; Cat# 7014) according to the manufacturer’s instructions. Briefly, samples underwent mechanical lysis via bead-based homogenization, followed by membrane binding, multiple wash steps, and elution via centrifugation. DNA quality and quantity were assessed using ThermoScientific™ NanoDrop™ UV-Vis Spectrophotometer (Thermo Fisher Scientific, Waltham, MA). Purified DNA was standardized to 20 ng/ul and transferred to the East Tennessee State University Molecular Biology Core Facility for library construction. Primers flanking the V3 and V4 domains of the bacterial 16S rRNA gene were used to amplify the DNA using the 16S Metagenomic Sequencing Library Preparation System (Illumina; San Diego, CA) following the manufacturer's instructions. Sequencing was performed by the University of Tennessee Genomics Center using the Illumina MiSeq machine. All sample sequencing was performed with the same flow cell.

***Microbiome analysis***

Sequence analysis was performed using the Qiagen CLC Genomics Workbench, version 23.0 (https://digitalinsights.qiagen .com), according to the manufacturer’s OTU clustering workflow. First, adapter sequences were trimmed, and the resulting samples were filtered for adequate sequencing coverage using the default workflow parameters, including a minimum of 100 reads per sample before OTU clustering. The Silva Complete database was used to cluster the high-coverage sequences into operational taxonomic units (OTUs) and OTUs with low abundance were removed prior to downstream analysis. Phylogenetic trees were constructed using the OTU data with MUSCLE alignment and used for alpha- and beta- diversity analysis. A permutational multivariate analysis of variance (PERMANOVA) was performed to determine overall diversity, and differential abundance analysis was used to identify taxa that differed among experimental groups.

***Short-chain fatty acid (SCFA) gas chromatography***

Approximately 500 mg of feces per sample was freeze-dried using a LABCONCO flask system. Samples were exposed to -50 °C for at least 12 hours at .077 mBar of pressure. Short chain fatty acids (SCFAs) were extracted using a procedure described previously^43^. In brief, SCFA extraction solution was prepared by adding oxalic acid (0.1 mol/L) and sodium azide (40 mmol/L) to 80 mg of freeze-dried fecal samples in 16 x 100 mm glass tubes, followed by vigorous mixing. Solution was centrifuged three times with supernatant removed after each centrifugation and stored in an ultralow freezer until use. SCFA concentrations were determined using a gas chromatography machine (Shimadzu GC2010) equipped with a capillary column (Sigma Aldrich, AB-Was Plus) with a flame ionization detector. The machine reported the concentration as parts per million (ppm) for each individual compound. SCFA concentrations were normalized to 80 mg of tissue for each sample. Values were subsequently converted and reported as nmol/mg.

***Statistical Analysis for SCFA***

Differential expression values for the microbiome 16S sequencing data were performed using CLC Workbench OTU clustering tutorial. All statistical analyses for SCFA were performed using GraphPad V10. SCFA concentrations were initially assessed for outliers using a ROUT test. Normality for the data sets for each unique SCFA was determined using Shapiro-Wilk and Kolmogorov-Smirnov tests.  A one-way ANOVA was used to assess differences among the three groups if the control group passed normality tests. A Kruskal-Wallis test was used if the control group was not normally distributed.

**Supplementary Figures**

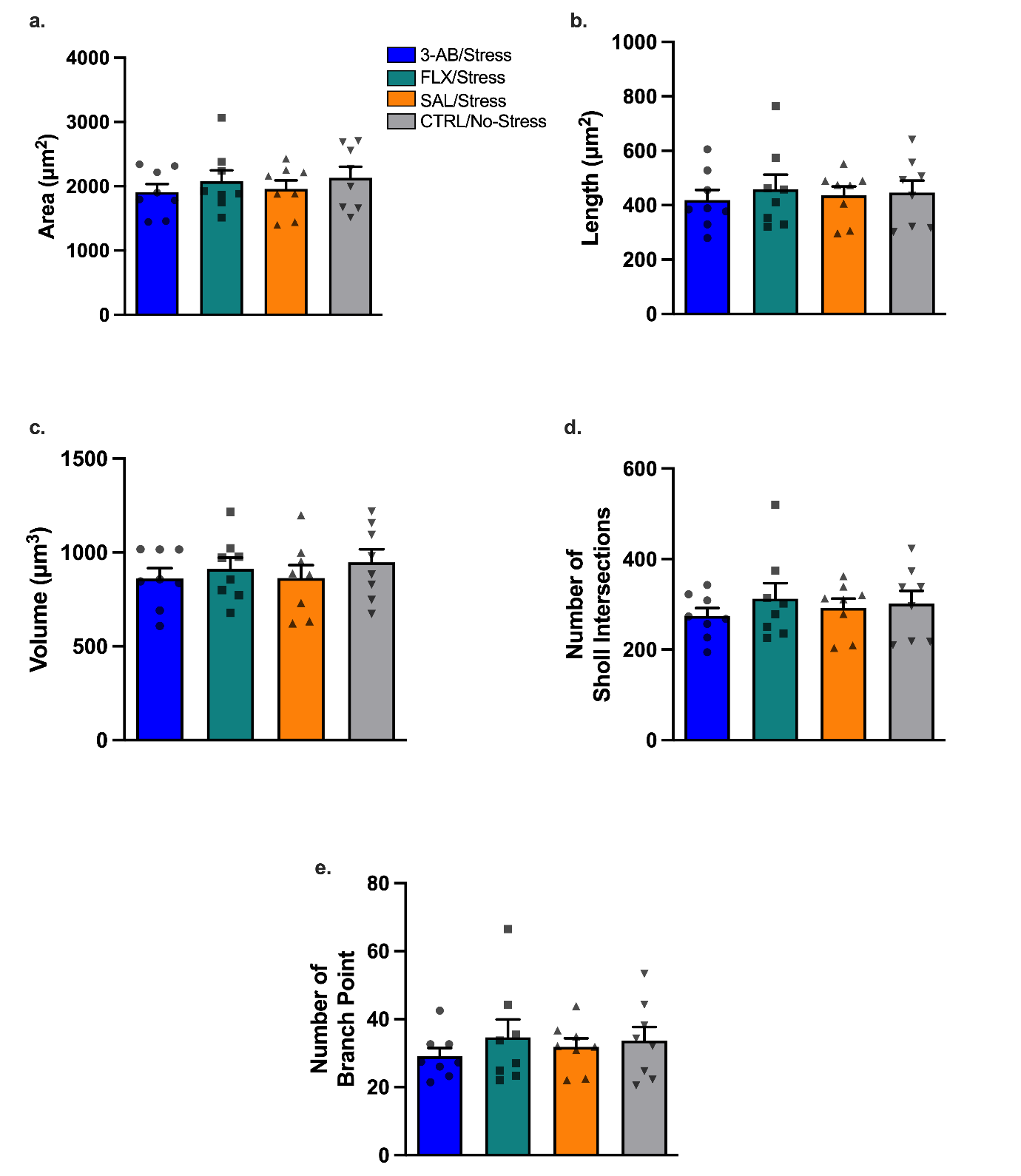

**Supplementary Figure S1. Microglial filament morphology in prefrontal cortex gray matter shows no treatment-related differences.** Microglial filament/ process characteristics measured from 3D reconstructed IBA-1-immunolabeled cells. (A) Filament area, (B) filament length, (C) filament volume, (D) number of sholl intersections, and (E) number of branch points. While stress significantly altered microglial soma morphology (Fig 7), no significant treatment effects were observed on filament parameters. Individual data points represent individual
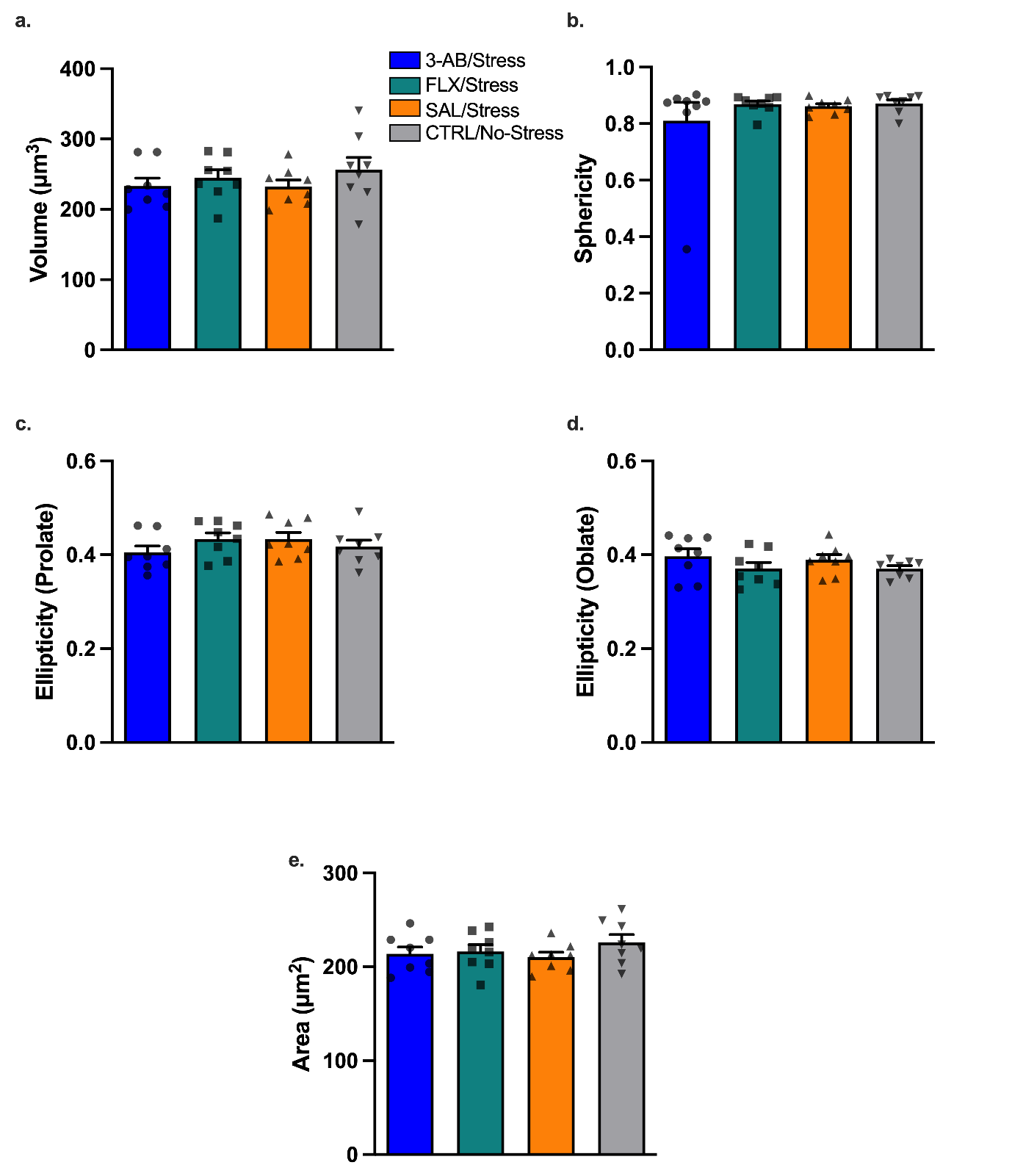
animals. Data are mean ± SEM (n = 8 per group).

**Supplemental Figure S2. Microglial soma morphology in prefrontal cortical white matter.** Microglial soma morphology parameters from 3D reconstruction of IBA-1-immunolabled microglia (15 cells per animal). (A) Soma volume, (B) soma sphericity, (C) soma ellipticity (prolate), (D) soma ellipticity (oblate), and (E) soma area. There were no significant differences in PFC white matter microglia soma morphology. Individual data points represent individual animals. Data are mean ± SEM (n = 8 per group).

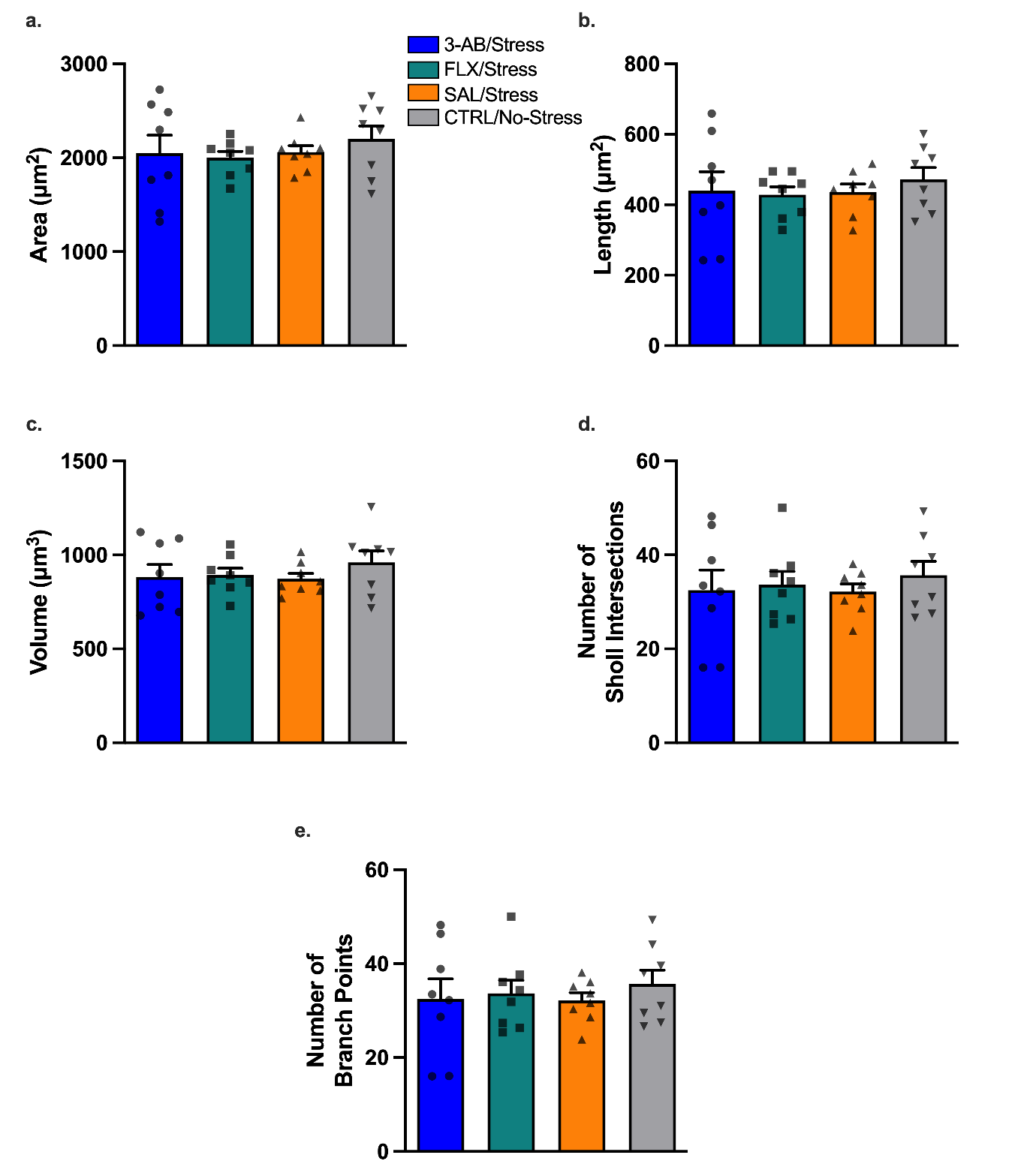

**Supplemental Figure S3. White matter microglial filament morphology in prefrontal cortex.** Microglial filament/ process characteristics measured from 3D reconstructed IBA-1-immunolabeled cells. (A) Filament area, (B) filament length, (C) filament volume, (D) number of sholl intersections, and (E) number of branch points. While stress significantly altered microglial soma morphology (Fig 7), no significant treatment effects were observed on filament parameters. Individual data points represent animals. Data are mean ± SEM (n = 8 per group).

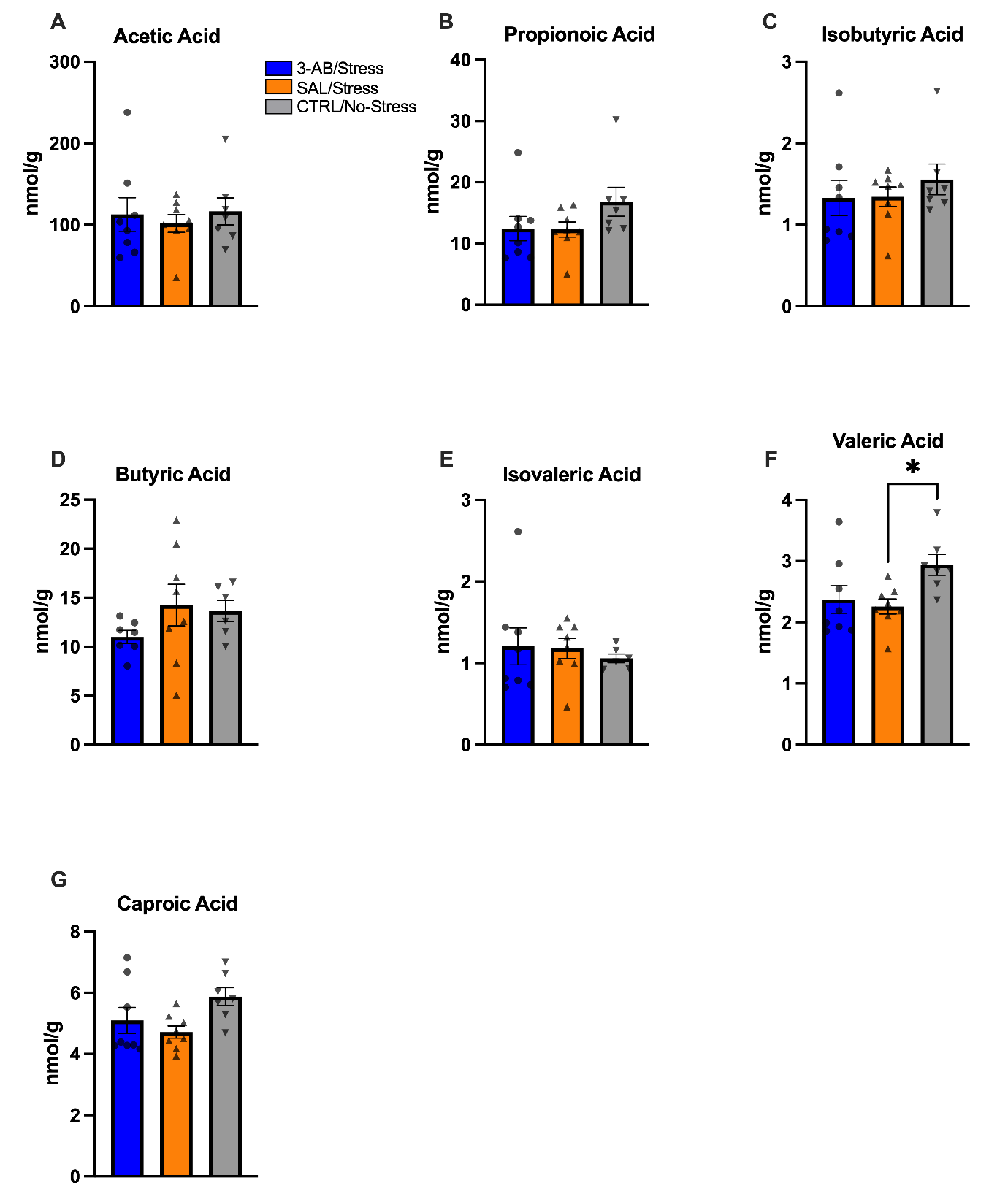

**Supplemental Figure S4. Chronic stress reduces cecal valeric acid concentrations.** Short-chain fatty acid (SCFA) concentrations measured by gas chromatography-mass spectrometry in cecal contents collected on day 12. (A) Acetic acid, (B) propionic acid, (C) isobutyric acid, (D) butyric acid, (E) isovaleric acid, (F) valeric acid, and (G) caproic acid. Rats in the SAL/Stress group showed significantly lower concentrations of valeric acid compared to the CTRL/No-Stress group (* p < 0.05), no significant treatment effects were observed in other SCFA. Individual data points represent individual animals. Data are mean ± SEM (n = 7-8 per group).

**Supplemental Table 1**. Differential abundance analysis for *Bacteroides dorei* AB242143.1.1490. Differential abundance analysis was performed using the QIAGEN CLC Genomics Workbench following OTU clustering with the SILVA 99 Complete database. Statistical significance was evaluated using false-discovery-rate (FDR) correction for multiple testing (Benjamin-Hochberg method).

| Comparison | Max Group mean | Fold Change | P-value | FDR | Bonferonni Adjusted P-value | Significance  FDR <.05 |
| --- | --- | --- | --- | --- | --- | --- |
| No Stress vs. Stress | 84.00 | -8.51 | 1.32 e-03 | 2.60 e-02 | 1 | Yes |
| Stress vs. Stress/3-AB | 84.00 | 9.95 | 5.77 e-04 | 3.13 e-02 | .50 | Yes |
| No Stress vs. Stress/3-AB | 10.25 | 1.17 | 8.17 e-01 | 9.39 e-01 | 1 | No |
